# The multidrug transporter Pdr5 from *Saccharomyces cerevisiae* is an ergosterol-modulated efflux pump

**DOI:** 10.64898/2026.10.29.762749

**Authors:** Lea-Marie Nentwig, Jakob Helmbrecht, Elise Kaplan, Sakshi Khosa, Haigang Song, Philipp Westhoff, Carol V. Robinson, Sander H. J. Smits, Andrzej Harris, Marten Exterkate, Ben F. Luisi, Lutz Schmitt

## Abstract

The pleiotropic drug resistance ABC transporter Pdr5 and its homologs play a critical role in multidrug resistance of numerous fungi, including pathogens. Previous studies demonstrated that alterations in the composition of the plasma membrane in which Pdr5 is embedded, specifically the ergosterol content, affected transport activity. Here we report the cryo-EM map of *Saccharomyces cerevisiae* Pdr5 in complex with ATP/ADP at 2.2 Å resolution, revealing densities on the surface of the transmembrane domains as well as inside the substrate-binding site, that were assigned as ergosterol, identified and quantified by GC-MS, and the sphingolipid M(IP)_2_C, identified via native MS. The ergosterol molecules are bound to the protein in well-defined grooves and unexpected locations.

We found that the activity of Pdr5 follows a Gaussian-like distribution, increasing with the ergosterol-concentration to reach a maximum before declining at higher concentrations, but this modulation was not observed in the presence of cholesterol, the sterol analogue found in mammalian cells. Mutational analysis corroborates the role of a single ergosterol-binding site in the modulation of Pdr5 activity.

Our structural analysis identifies a protein-lipid network that could serve as a center for lipid engagement linking the surrounding lipid environment to the substrate-binding region of Pdr5.

These findings advance our understanding of Pdr5 function by demonstrating that the lipid environment is closely linked to Pdr5 structure and activity.

## Introduction

Pathogenic fungal infections are a growing risk to human health ^1^, but our arsenal to combat these infections is restricted to five major antifungal drug classes ^2^. Since their introduction in 1980, azoles have been the primary drug class used for the treatment of fungal infections ^3^ by targeting the ergosterol biosynthesis pathway. Ergosterol is the major sterol in fungal plasma membranes, where it contributes to membrane stability by regulating membrane fluidity and rigidity. Its role is analogous to that of cholesterol in mammalian cells. Consequently, the similarity between fungal ergosterol and mammalian cholesterol contributes to the challenge of developing selective antifungal therapies, as interactions with cholesterol can lead to host toxicity^4^. At the same time, the overuse of azoles in therapy has led to the development of a number of resistant fungi, some of which are multidrug-resistant ^3^. Especially for azoles, the major resistance mechanism is active drug efflux mediated by membrane transport proteins of the ABC transporter superfamily ^1^.

Functional ABC transporters share a common architecture consisting of two nucleotide-binding domains (NBDs) and two transmembrane domains (TMDs). The NBDs bind and hydrolyze ATP to provide the energy for transport, while the TMDs are located in the plasma membrane which mediate substrate transport across the lipid bilayer ^5^. In the case of azole resistance in human pathogenic fungi, the PDR (pleiotropic drug resistance) subfamily plays an important role ^6^.

A characteristic feature of the PDR proteins is the asymmetry in their NBDs originating from deviations in the characteristic motifs necessary for ATP binding and hydrolysis within the ABC transporter family ^7^. This asymmetry leaves one of the two nucleotide-binding sites (NBSs) degenerated or deviant, catalytically inactive and unable to hydrolyze ATP ^8^. Pdr5 from *Saccharomyces cerevisiae* is the founding member of this ABCG subfamily and presents one of the most severe degree of NBD asymmetry ^9^. This protein was identified more than three decades ago and studied extensively ever since ^7^. The establishment of a purification protocol to isolate Pdr5 in a functional state as well as advances in cryogenic-electron microscopy (cryo-EM) of membrane proteins, enabled the elucidation of the structures of Pdr5 ^10^. Subsequently, structures of Pdr5 homologs Cdr1 and Cdr2 from the *Candida* species have been reported ^11, 12, 13, 14, 15^. Organisms of the *Candida* species such as *C. albicans*, *C. auris*, or *C. glabrata* that have been categorized in the ‘critical priority list’ of drug resistant fungi by the World Health Organization (WHO) in 2021 ^16^.

The current structures of Pdr5 in the presence or absence of nucleotide and/or substrate form the foundation of the proposed transport cycle mechanism, including drug translocation and ATP hydrolysis ^10^. Additionally, a linker domain located between the two halves of the protein functions as a nucleotide sensor that recognizes binding of the nucleotide in the deviant NBS and dictates ATP hydrolysis in the catalytically active NBS ^10^.

As stated above, the fungal plasma membrane contains ergosterol which is synthesized through a tightly regulated, ATP-demanding biosynthetic pathway, reflecting its essential role in the fungal plasma membrane ^17, 18^. In *S. cerevisiae*, ergosterol is tightly packed together with sphingolipids in membrane microdomains. Likewise, clustering of Pdr5 into membrane microdomains was demonstrated ^19^, possibly providing an optimal environment for its function ^20, 21^. The tight packing of ergosterol and sphingolipids not only accommodates integral membrane proteins but is also associated with drug susceptibility ^22^. Importantly, lipids and sterols not only influence the physical properties of the membrane but are also directly involved in membrane protein function through specific protein-lipid interactions ^17^. Two prominent examples of cholesterol-dependent activity are the human ABC transporters ABCB1 and ABCG2. Their ATPase activity, substrate transport and conformational dynamics are modulated by cholesterol, highlighting the importance of the membrane environment for transporter function ^23, 24, 25^. To what extent similar regulatory mechanisms apply to the yeast ABC transporter Pdr5 remains unclear. Hence, understanding the relationship between Pdr5 and its local membrane environment, *i.e.*, possible interactions with membrane constituents, is required to gain detailed insights into the regulation of this transporter, ultimately advancing our understanding of multidrug efflux pumps in yeast.

Here, we report the cryo-EM map of ATP/ADP bound Pdr5 in the inward-facing (IF) conformation at 2.2 Å. In this map, densities identified as co-purified lipids and ergosterol molecules were observed. Additionally, the most prominent sphingolipid of yeast ^26^, mannosyl-di-(inositolphosphoryl)-ceramide (M(IP)_2_C) occupies the substrate-binding site (M(IP)_2_C^ligand^) and is localized in close vicinity of the substrate exit channel of Pdr5 (M(IP)_2_C^OL^). A detailed functional and mutational analysis revealed that ergosterol is necessary and sufficient to modulate the ATPase activity of Pdr5. These results emphasize the importance of ergosterol as prime regulator of Pdr5 and suggest a cross-talk ^27^ between ergosterol biosynthesis and drug efflux activity.

## Results

### A high-resolution cryo-EM map of Pdr5 reveals a local lipid environment

Structures of Pdr5 have been reported for the apo-state (pdb entry 7P03), with bound nucleotides (ATP and ADP; pdb entry 7P04), in the presence of ATP and sodium-orthovanadate (pdb entry 7P06), trapping the protein in the outward-facing state as well as in the presence of ATP and the transport substrate rhodamine 6G (pdb entry 7P05). These structures are in the resolution range from 2.9–3.8 Å ^10^.

Improvements in cryo-EM data collection and processing, as well as in the purification protocol of Pdr5 in the detergent PCC-α-M ^28^ now led to an increase in resolution to 2.2 Å of the ATP/ADP bound Pdr5 cryo-EM map. An ATP molecule is present in the deviant NBS1 and ADP/Mg^2+^ were found in the catalytically active NBS2, in agreement with the previously published structure. The nature of the bound nucleotides as well as the cofactor and coordinating water molecules could be unambiguously identified in the electron density map (Fig. S1).

Although the overall architecture of the protein is consistent with the previously reported Pdr5 structure (root mean square deviation (rmsd) of 0.8 Å across 1353 C_α_ atoms for the ATP/ADP bound structure), the 2.2 Å reconstruction provides improved map quality (Table S1), with well-resolved density for the transmembrane helices throughout the TMDs (Fig. S2). Additionally, we now observe EM density for water molecules located in the NBDs, the extracellular domains (ECDs), and the putative position of the phospholipid headgroups. We also observe additional density for molecules surrounding the protein in regions corresponding to the inner and outer leaflets of the plasma membrane (Fig S3). Furthermore, non-protein density can be observed in the location of the substrate binding site, although only Mg^2+^ and ATP, present in the NBDs, was added to the sample (Fig. S4).

These additional densities were assigned to two molecules of N-acetyl glucosamine (NAG) covalently linked to residue 734, to phospholipids with one phosphatidylinositol-4,5-bisphosphate (PIP_2_), one phosphatidylglycerol (PG), and two phosphatidylethanolamine (PE) molecules, and to ergosterol (four molecules), the sterol-component of the plasma membranes of fungi. Especially for ergosterol, the quality of the density allowed an unambiguous placement (Figure S4). However, two densities, one in the ligand-binding site and one in proximity of the substrate exit channel, could not be easily assigned. To guide their identification, we performed native MS analysis (see paragraph below).

### Identification of sphingolipids bound to Pdr5

While the densities corresponding to ergosterol and several phospholipids (PIP_2_, PG, PE) could be readily assigned in the high-resolution cryo-EM map of Pdr5, two additional densities remained unassigned. We therefore performed native mass spectrometry (MS), which independently identified a co-purified lipid species that matched this density and thus, enabled the assignment (Fig. S5).

Pdr5 is N-glycosylated at N734 and phosphorylated, which increases the complexity of the pattern of peaks in the native mass spectra. However, upon phosphatase treatment of Pdr5 N734A variant, native mass spectra revealed three major lipids adducts with mass shifts of +834 Da, +1,371 Da and +2,199 Da (Fig. S5a-b). The +836 Da is consistent with phosphatidylinositol (PI 34:1), whereas the +1,371 Da could not be assigned to canonical lipids. Pseudo-MS^2^ analysis under high in-source activation conditions released a series of ligands with *m/z* 1,326.7843, 1,342.7800, 1,358.7751, 1,370.8107, and 1,386.8061 (Fig. S5c) at low mass range. Isolation and subsequent fragmentation of ion (pseudo-MS^3^) at *m/z* 1,358.7751 identified it as M(IP)_2_C 44:0;O4 (Fig. 5d) which was modelled as the representative sphingolipid in the cryo-EM structure. These two densities represent the headgroup and parts of the acyl chains of mannosyl-di-(inositol-phosphoryl)-ceramide (M(IP)_2_C) ^26^, a complex and highly abundant sphingolipid in the plasma membranes of fungi (Fig. 1, Fig. S4). While the density present in the drug-binding site of Pdr5 accommodates the mannosyl-di-(inositol-phosphoryl)-headgroup, no reliable density can be observed for most of the lipid tails, presumably due to their flexibility within the detergent micelle (Fig. 2 and Fig. S4). Focusing on the M(IP)_2_C^ligand^ headgroup, a comparison with the rhodamine 6G (R6G) bound structure of Pdr5 ^10^, reveals that this M(IP)_2_C occupies the ligand-binding site of Pdr5 (Fig. S6) and most of the interacting residues are identical for the two ligands.

**Figure 1.**
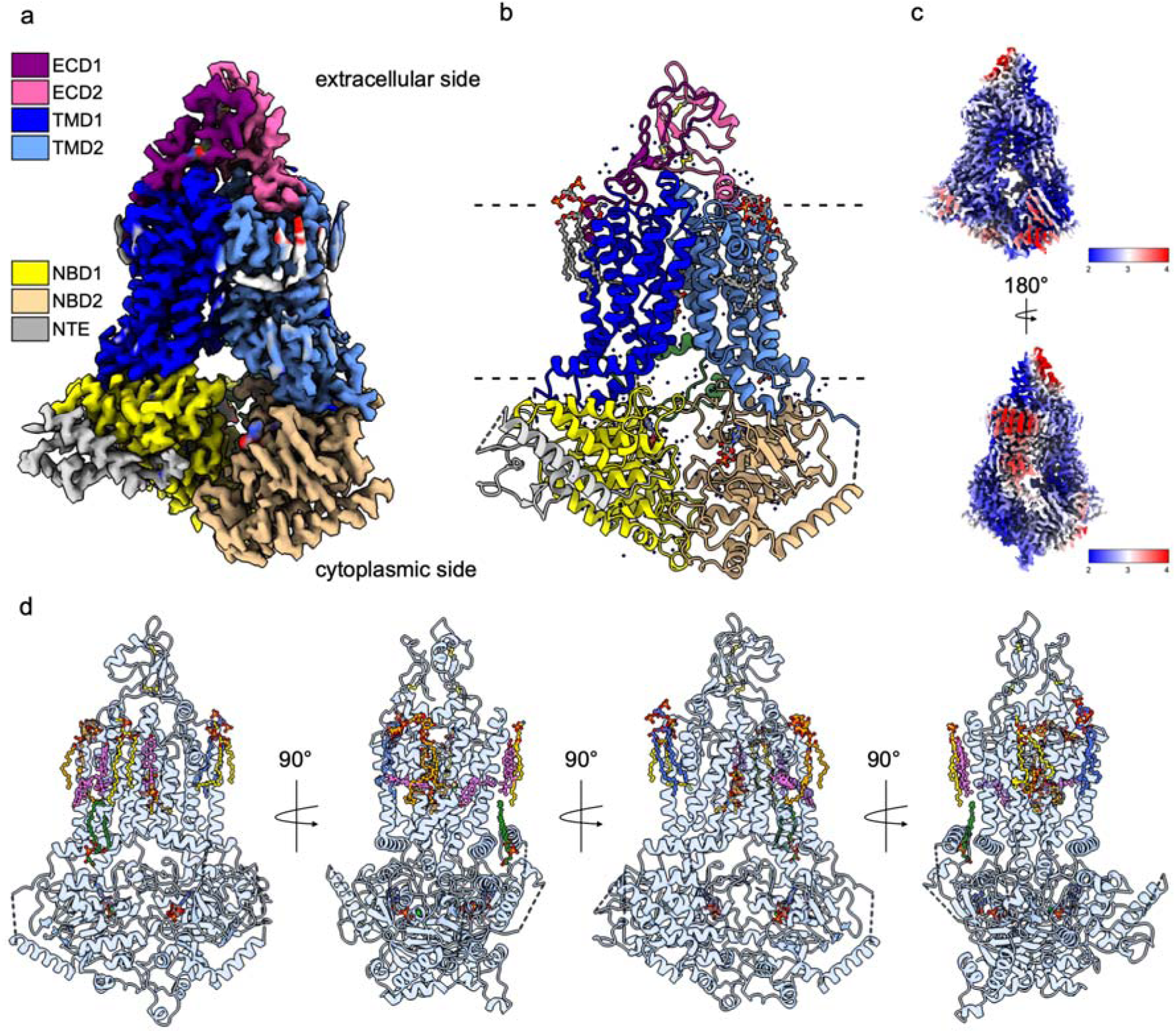
Cryo-EM structure of Pdr5. (a) Cryo-EM density map, color-coded as follows: N-terminal extension (grey, residues 1– 127), NBD1 (yellow, residues 128– 140 and 169– 490), TMD1 (blue, residues 491– 716 and 774-795), ECD1 (magenta, residues 717– 773), nucleotide sensor (green, residues 141– 168 and 796– 861), NBD2 (wheat, residues 861– 1166), TMD2 (light blue, 1167– 1404 and 1471– 1511), ECD2 (light magenta, residues 1405– 1470). (b) Overall structure of Pdr5 in complex with ATP in NBS1 (deviant site) and ADP/Mg^2+^ in NBS2 (active site). Nucleotides and bound ligands are shown in ball-and-stick representation. Water molecules and Mg^2+^ are depicted as spheres (blue for water and green for Mg^2+^). The three disulfide bridges of the ECDs (residues 722/742, 1411/1455, and 1427/1452) are shown in yellow ball-and-stick representation. (c) Local resolution of the cryo-EM density map, as calculated by chimeraX ^29^. The color scale indicates the local resolution in Å. (d) 90° rotations of Pdr5 in the plane of the membrane. To highlight the bound ligands, nucleotides, lipids, and ergosterol, Pdr5 is shown in a transparent, blue cartoon representation. Color-coding of the ligands in ball-and-stick representation is as follows: PIP_2_ in blue, PG lipids in green, the sphingolipids M(IP)_2_C in orange, PE lipid in yellow, ATP and ADP in element color coding, and ergosterol in magenta.

**Figure 2.**
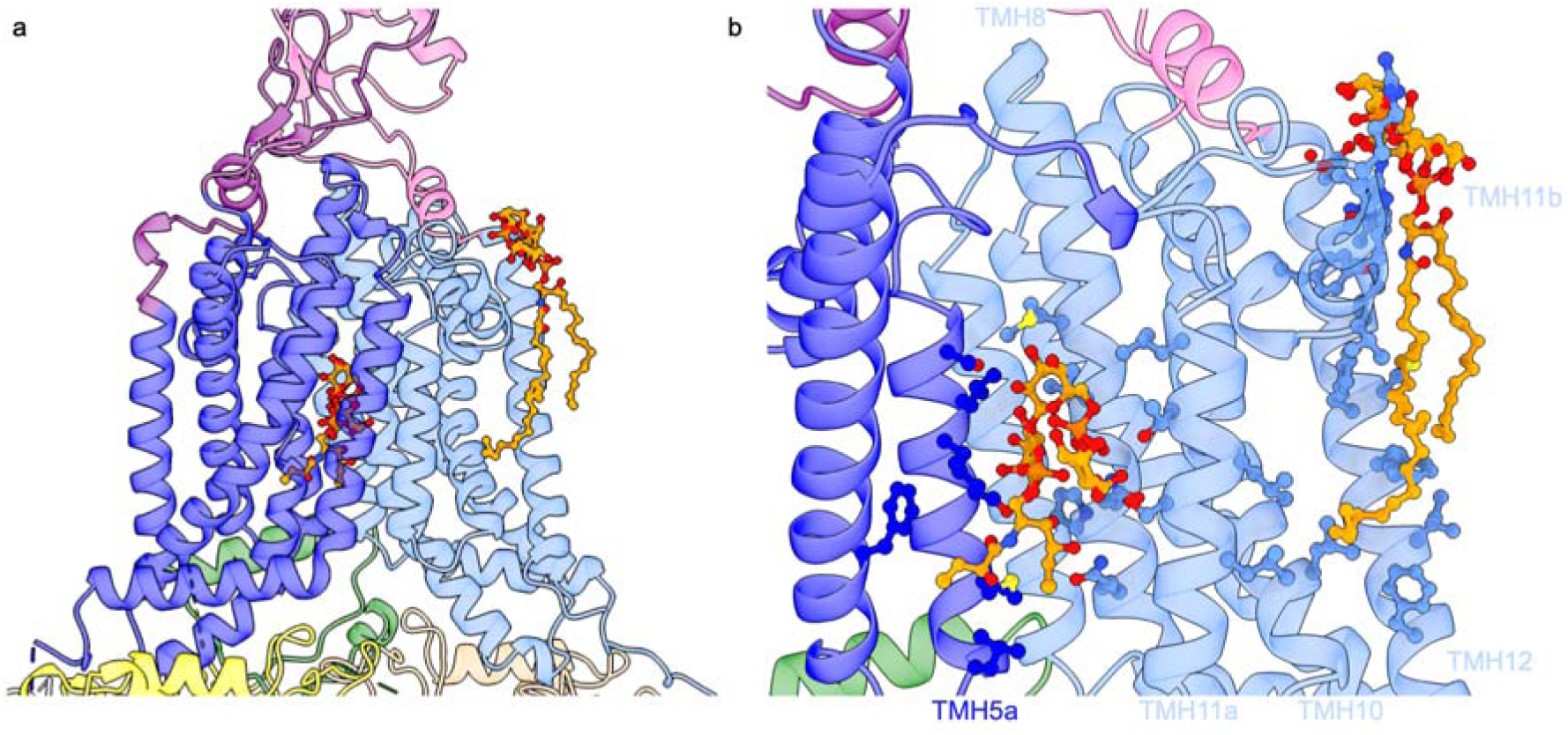
Interactions of Pdr5 with the sphingolipid M(IP)_2_C. (a) Binding region of the two M(IP)_2_C molecules shown in orange ball-and-stick representation (oxygen atoms are shown in red). Color-coding is as in Fig. 1a. (b) Close-up view of the M(IP)_2_C interactions with Pdr5. For clarity, TMH1b, TMH2, and TMH3 are not shown. Helices involved in the stabilization of the two M(IP)_2_C molecules are labeled. Residues within 4 Å of the M(IP)_2_C^ligand^ molecule in the substrate-binding site are located in TMH2 (top to bottom): L565, F566, L572, I575; TMH4: F653; TMH5a (bottom to top): L663, M667, S671, L674, L675, S678; TMH8 (top to bottom): M1243, V1246, N1249, P1250, Q1253; TMH11a (bottom to top): S1360, T1364, L1367. Residues within 4 Å of the M(IP)2C^OL^ on the surface of Pdr5 are located in TMH10: L1344; TMH11a (bottom to top): L1358, L1362; TMH11b (top to bottom): R1381, I1384, R1388, V1389; TMH11c: Y1394; TMH12 (top to bottom)): E1473, R1476, I1480, C1483, Y1484, A1486, F1487, I1490, F1495.

A second density outside TMH12, M(IP)_2_C^OL^, which is part of the exit gate of Pdr5, is compatible not only with the M(IP)_2_C-headgroup, but also larger parts of the ceramide moiety, indicating less flexibility (Fig. 2, Fig. S4).

### Overexpressed Pdr5 is present in lipid microdomains *in vitro*

Ergosterol and M(IP)_2_C 44:0;O4 are the most abundant lipid species in yeast ^30^. Our observation that Pdr5 co-purifies with both ergosterol and M(IP)_2_C, suggests the transporter may be associated with sterol- and sphingolipid-rich membrane microdomains even under conditions of overexpression. At the endogenous level, a clustering of Pdr5 in microdomains was already demonstrated ^19^. To investigate whether overexpression might induce mis-localization, detergent-resistant microdomains (DRMs) were isolated via density-gradient centrifugation and microdomain association of overexpressed Pdr5 was assessed via Western blot analysis. Fractionation revealed co-migration of Pdr5 with the marker protein Pma1 in DRM fractions, which contains different types of plasma membrane microdomains (Fig. 3a). Isolation of intact DRM fractions was confirmed by LC-MS analysis, revealing the DRM marker lipids mannosyl-inositolphosphoryl-ceramide (MIPC) and inositolphosphoryl-ceramide (IPC) in the low-density fractions 2 and 3 (Fig. 3b, c). Hence, Pdr5 is indeed microdomain-associated, even under conditions of overexpression, but is not required for the formation of microdomains.

**Figure 3.**
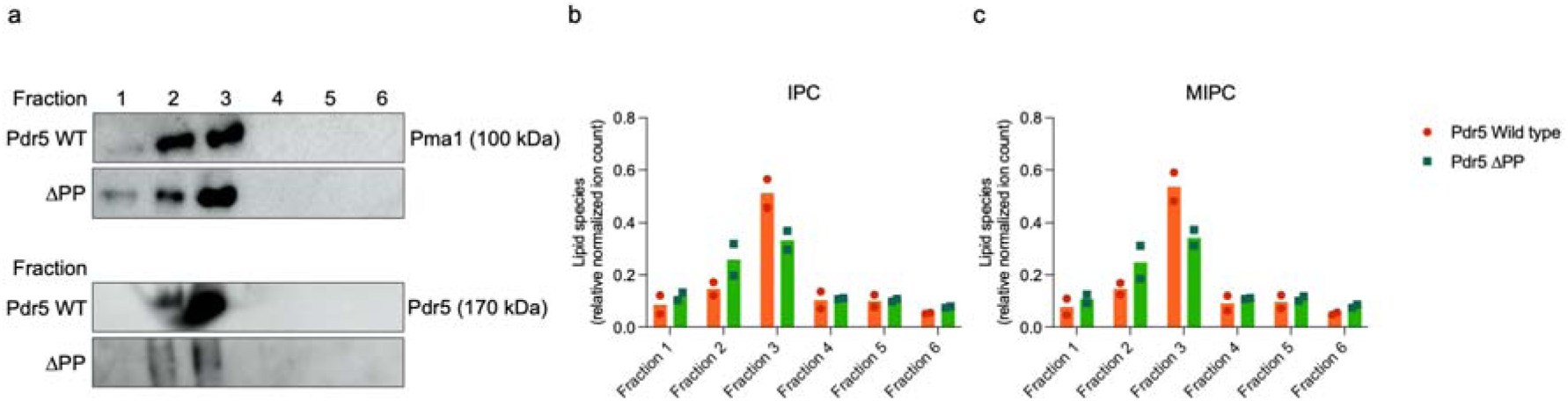
Pdr5 is associated with membrane microdomains. (a) DRMs were isolated from yeast cells either expressing Pdr5 Wild type (Pdr5 WT) or the knock-out strain (ΔPP). Isolated fractions were resolved by SDS-PAGE and detected by Western Blot using an antibody against the detergent-resistant microdomain (DRM) marker Pma1 or Pdr5. LC-MS analysis of DRM lipid markers IPC (b) and MIPC (c). Relative signal intensities of each fraction were calculated as the percentage of the total signal intensity across all fractions for two independent experiments.

### Pdr5 engages avidly four ergosterol molecules

The four molecules of ergosterol, identified unambiguously in the density, are bound to the surface of TMD2 within distinct grooves (Fig. 4), which differs from the location of cholesterol molecules found in ABCG2 ^31^ (Fig. S7). Beginning with the ergosterol closest to the substrate-binding site near the center of the transporter, the remaining sterols are described in anti-clockwise order along TMD2.

**Figure 4.**
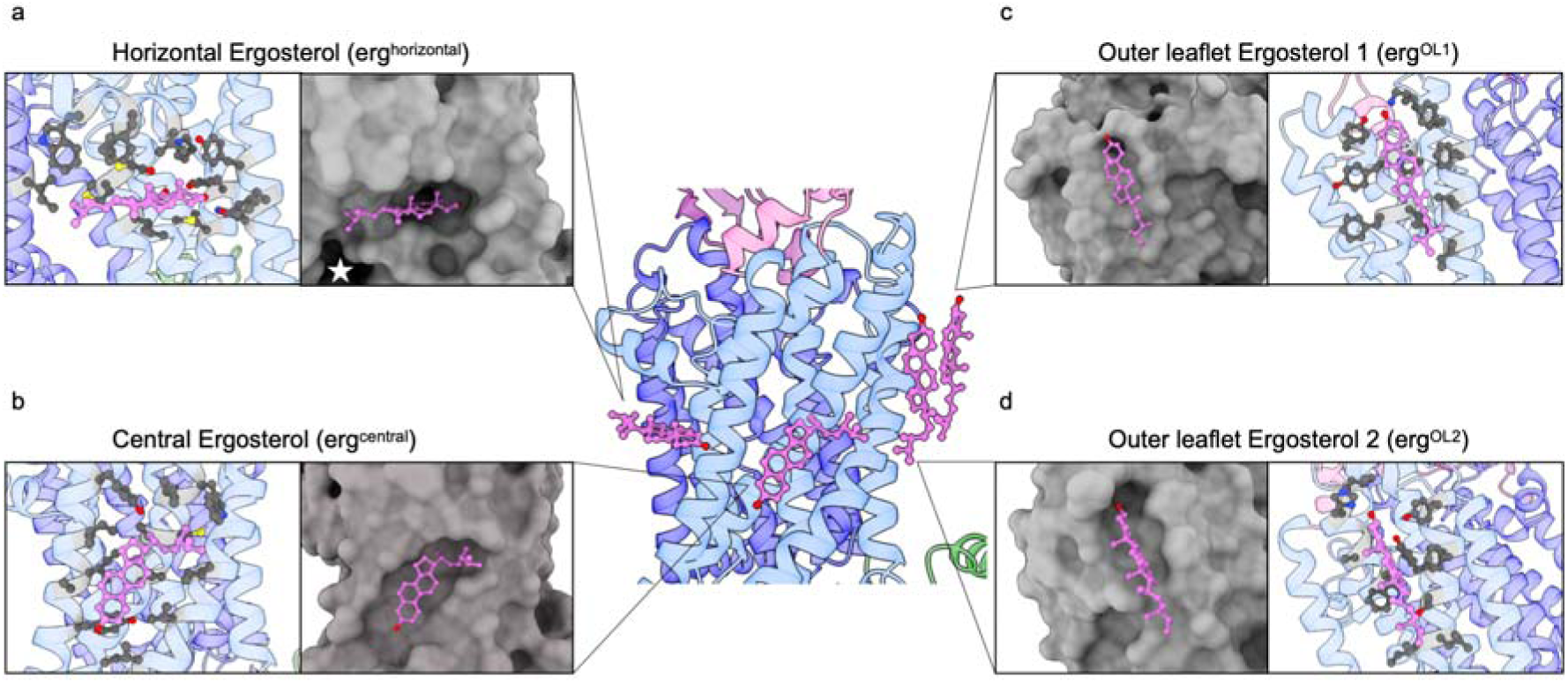
The ergosterol binding sites of Pdr5. The central panel shows the four ergosterols bound to Pdr5. For simplicity, all other ligands were omitted and Pdr5 is shown in transparent cartoon representation. Color-coding is as in Fig. 1a. Detailed view of the ergosterol-binding cavity and the interacting protein residues for erg^horizontal^ (a), erg^central^ (b), erg^OL1^ (c) and erg^OL2^ (d). Residues within 4 Å of the bound ergosterols are shown in ball-and-stick representation.

The ergosterol molecule, referred to as erg^horizontal^ (Fig. 4a), is found in a horizontal orientation at the center of the membrane, between the two phospholipid leaflets, accommodated within a deep groove of the protein, highlighting a specific interaction with Pdr5. Notably, erg^horizontal^ is positioned in close proximity of the substrate-binding entry site, which is located in the center of the protein (marked with a white star in Fig. 4a).

The ergosterol molecule, referred to as erg^central^ (Fig. 4b), also occupies a deep groove at the protein surface, but adopts a tilted conformation with respect to the membrane plane. While the sterol ring system is positioned closer to the inner leaflet of the lipid bilayer, the hydroxyl headgroup is rotated away for the rings and extends towards the middle of the phospholipid bilayer. Interestingly, no specific protein interaction shielding the hydrophilic OH-group was detected.

The two other ergosterol molecules, erg^OL1^ (Fig. 4c) and erg^OL2^ (Fig. 4d), are located at the site of Pdr5 that will be embedded into the outer leaflet of the phospholipid bilayer. Both molecules are bound within less well-defined grooves in the protein. One has to note that, in contrast to mammalian cells, 80 % of the ergosterol molecules in the plasma membrane of yeast localize to the inner leaflet ^32, 33^, whereas these two ergosterol molecules are positioned at the water-lipid interface of the membrane outer leaflet.

The number of ergosterol molecules per Pdr5 estimated by gas chromatography-mass spectrometry (GC-MS) is 4.0 ± 0.3 which is in agreement with the observation from the cryo-EM map (Fig. S8b).

### The ATPase activity of Pdr5 is ergosterol-modulated

The purification of functional Pdr5 critically depends on the selection of a suitable detergent ^34^. So far, only PCC-α-M retained ATPase activity of purified Pdr5 ^34^. The presence of ergosterol molecules in Pdr5 solubilized and purified in PCC-α-M raised the question whether different detergents differ in their ability to preserve the protein-associated ergosterol during solubilization and purification. DDM has been reported to solubilize Pdr5 ^35^ and was used to determine its negative stain EM structure ^36^. We therefore purified Pdr5 using either PCC-α-M or DDM and subsequently determined ATPase activity of the detergent-solubilized protein as well as quantified the number of co-purified ergosterol by GC-MS, respectively. Pdr5 in DDM did not show ATPase activity, whereas Pdr5 purified in PCC-α-M was active (Fig. S8a) ^34^. Indeed, GC-MS quantification revealed 0.30 ± 0.01 ergosterol molecules co-purified per Pdr5 molecule in DDM (Fig. S8b), whereas 4.0 ± 0.3 ergosterol molecules were detected in PCC-α-M purified Pdr5, suggesting that the presence of ergosterol is essential for Pdr5’s ATPase activity.

To further address the functional role of ergosterol, we analyzed the influence of varying amounts of sterols on Pdr5 ATPase activity via a previously described *in vitro* ATPase assay ^37^. In short, Pdr5 purified in PCC-α-M was reconstituted into liposomes containing varying amounts of ergosterol (5–20 % (w/w), while the remaining lipids were POPC, POPE, POPS (3:3:2 (w/w)). To ensure that any observed effect can be attributed specifically to ergosterol, we included cholesterol, the mammalian counterpart to ergosterol, as a control in our analysis. To validate cholesterol as a suitable mimic, the membrane fluidity, order and lipid packing were investigated via measurement of anisotropy and general polarization of the liposomes (Fig. S9). As evident from the data, ergosterol and cholesterol modulated the physico-chemical parameters of the bilayer similarly, confirming that cholesterol indeed serves as a mimic for ergosterol and is not changing the overall membrane properties. We therefore proceeded with the reconstitution of purified Pdr5 into these liposomes ^37^ containing either 5, 10, 15 or 20 % (w/w) of ergosterol or cholesterol. Surprisingly reconstitution of Pdr5 into liposomes lacking a sterol, was not successful. To calculate the Pdr5-specific ATPase activity, the efficiency of reconstitution and the orientation of the transporter in the liposomes with respect to the fraction of NBDs accessible to ATP for hydrolysis were considered ^37^ (Fig. S10).

The results revealed a clear picture: the ATPase activity of Pdr5 scales with increasing ergosterol content, reaching its maximum activity of 642 ± 39 nmol/(mg*min) at 15 % (w/w) ergosterol. Further increasing the ergosterol concentration up to 20 % (w/w), resulted in a decrease in ATPase activity of Pdr5 to 420 ± 38 nmol/(mg*min) (Fig. 5a, Table S2). The ATP hydrolysis-deficient control, Pdr5 E1036Q (Pdr5 EQ) did not display any significant ATPase activity, confirming that the measured ATPase activity is indeed Pdr5 specific. In contrast to ergosterol, the ATPase activity of Pdr5 in liposomes with varying amounts of cholesterol (5-20 % (w/w)), does not change, although the overall activity seems somewhat higher in comparison to ergosterol-containing proteoliposomes with a *V*_max_ value of around 1000 nmol/(mg*min) (Fig. 5b, Table S2).

**Figure 5.**
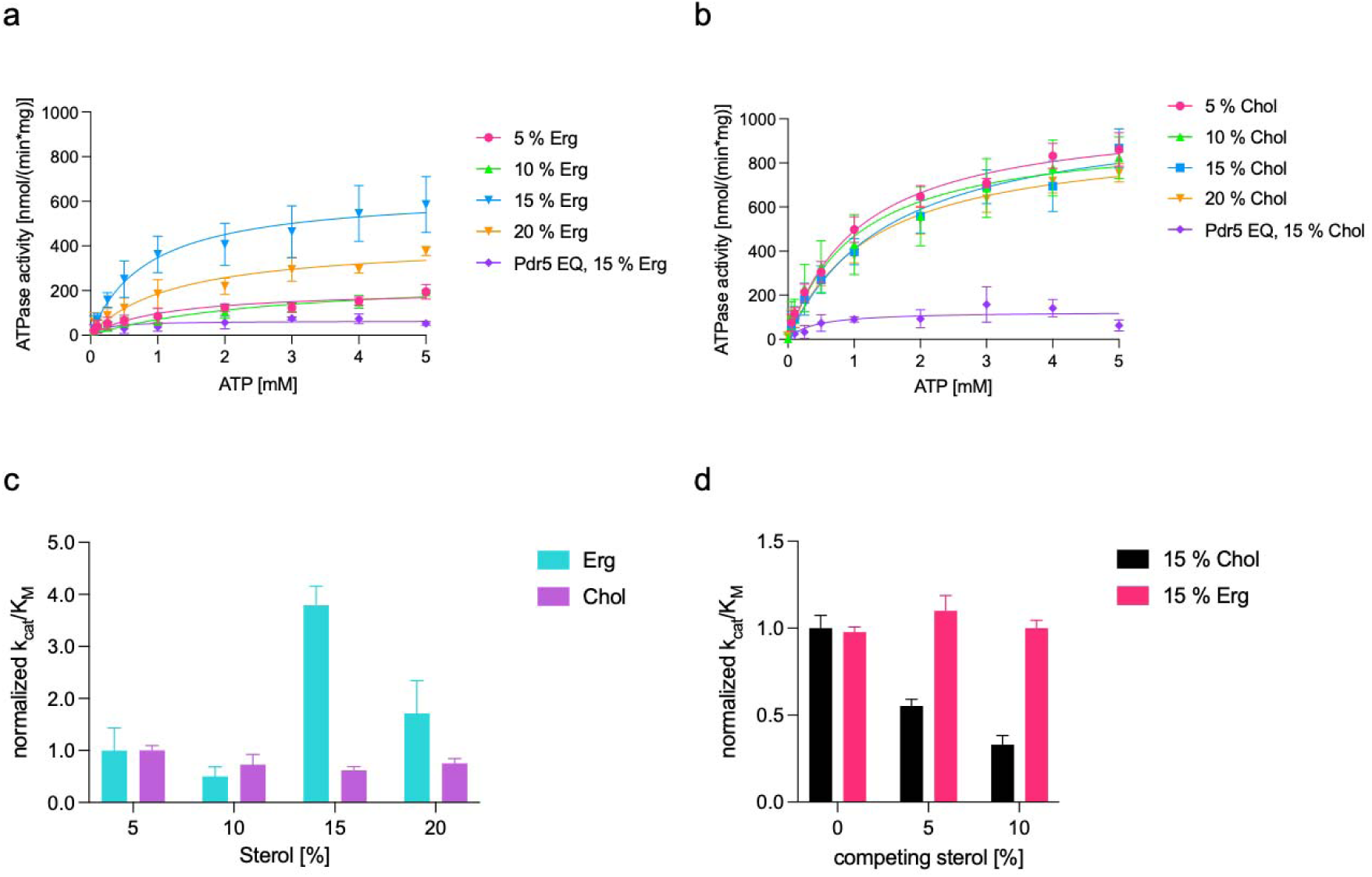
ATPase activity of Pdr5 WT reconstituted in synthetic liposomes. Activity in the presence of varying amounts of ergosterol (a) and cholesterol (b) and corresponding catalytic efficiency normalized to either 5 % (w/w) ergosterol or cholesterol, respectively (c). (d) Catalytic efficiency of the ATPase activity of Pdr5 WT in synthetic liposomes containing 15 % (w/w) cholesterol with the addition of 5 or 10 % (w/w) ergosterol normalized to 15 % (w/w) ergosterol or cholesterol in the absence or presence of competing sterol. Error bars represent the SD of at least three independent experiments.

To account for potential compensation between *V*_max_ and *K*_M_, we determined the catalytic efficiency (*k*_cat_/*K*_M_) of Pdr5 at each sterol concentration (Fig. 5c, Table S2). This strengthened the observed trend, with a catalytic efficiency remaining low for Pdr5 reconstituted into liposomes containing 5 and 10 % (w/w) ergosterol, increasing to a maximum at 15 % (w/w), and subsequently decreasing at 20 % (w/w) ergosterol. Hence, the ATPase activity of Pdr5 is modulated in an ergosterol-dependent manner, with behavior that indicates complex allosteric effects.

To further investigate this modulating effect and probe the sterol selectivity of the transporter, we performed ATPase activity assays of Pdr5 reconstituted into liposomes with a combination of cholesterol (15 % (w/w)) and varying amounts of ergosterol (0, 5 or 10 % (w/w)) resulting in liposomes with a total sterol content of 15, 20 or 25 % (w/w), respectively. Remarkably, in the presence of 15 % (w/w) cholesterol, the addition of ergosterol leads to a reduction of ATPase activity towards levels observed for Pdr5 WT reconstituted into liposomes with ergosterol alone (Fig. 5d), confirming that the modulating effect on Pdr5 is restored by the addition of ergosterol (Fig. 5d, Table S2). Vice versa, the catalytic activity of Pdr5 in liposomes containing 15 % (w/w) ergosterol is not influenced by additional cholesterol (Fig. 5d, Table S2). Thus, the ATPase activity of Pdr5 is governed by ergosterol whenever it is present, irrespective of the cholesterol concentration, whereas cholesterol alone cannot maintain the elevated activity once ergosterol is introduced. This trend is also represented by the catalytic efficiency (Fig. 5d).

### The central ergosterol (erg^central^) modulates the ATPase activity of Pdr5

We sought to identify the ergosterol molecule primarily responsible for the modulating effect on the ATPase activity of Pdr5. For this purpose, we applied a similar approach as described for Upc2 ^38^, a fungal transcription factor which is able to sense the cellular level of ergosterol, thereby regulating its biosynthesis. Modeling of cholesterol in the ergosterol-binding pocket sterically clashes with the latter, explaining the inability of this transcription factor to bind cholesterol. Likewise, we modeled cholesterol into the erg^central^ binding pocket of Pdr5. As shown in Fig. 6a, there is a good fit between the shape of the cavity and the ergosterol molecule. In contrast, clashes were observed for leucine at position 1329 as well as tyrosine at position 1478 (Fig. 6b, highlighted by a black circle) when cholesterol is docked into the groove (Fig. S11). We hypothesized that the substitution of both amino acid residues to alanine should result in a Pdr5 variant that can better accommodate cholesterol. Functional analysis of this Pdr5 mutant (L1329A and Y1478A, referred to as Pdr5 L/A–Y/A) reconstituted into liposomes containing varying amounts of cholesterol as previously described revealed a modulatory effect on ATPase activity (Fig. 6c), as seen for the Pdr5 WT and ergosterol (Fig. 5a, Table S2). This strongly suggests that the ergosterol-mediated modulation of Pdr5 takes place in the central binding pocket. This modulating effect was not observed for the Pdr5 WT reconstituted into varying amounts of cholesterol (Fig 5b, Table S2).

**Figure 6.**
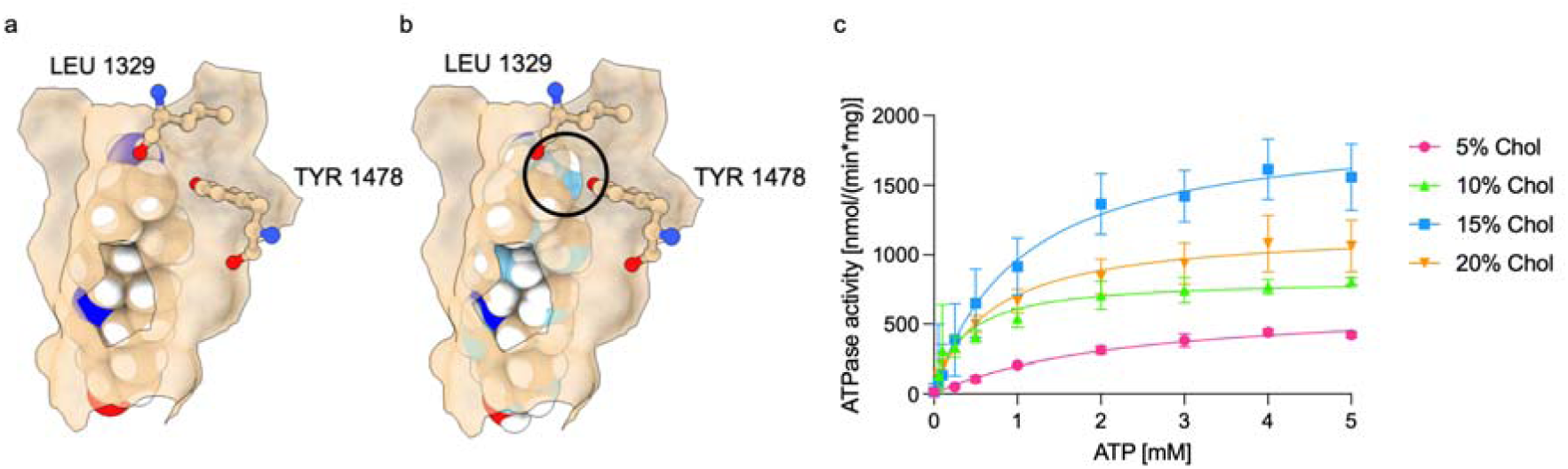
Structural basis for specificity for ergosterol over cholesterol in Pdr5. (a) Binding cavity of the central ergosterol (erg^central^). Erg^central^ is shown in light-brown spheres with hydrogens in white and the atoms which differ to cholesterol (methyl group at C24 and double bond between C22 and C23, Fig. S11) in blue. (b) Cholesterol (light blue sphere and hydrogens in white) was modelled in the erg^central^ location. Steric clash between the wall of the binding cavity is highlighted by a black circle. The amino acid residues inducing this steric clash are labelled (L1329 and Y1478). (c) ATPase activity of Pdr5 L/A–Y/A reconstituted in synthetic liposomes in the presence of varying amounts of cholesterol. Error bars represent the SD of at least three independent experiments.

In the presence of ergosterol, the mutant Pdr5 L/A–Y/A displayed the same modulation of ATPase activity as the wild type (Fig. S12). We did not analyze the binding sites of the two outer leaflet ergosterols as the shape complementarity was not as pronounced as that of the central groove. In the case of erg^horizontal^, small clashes were observed when cholesterol was docked into the structure.

### Helices TMH11 and TMH12 serve as a connecting hub for ergosterol and M(IP)_2_C

The clustering of erg^horizontal^ and erg^central^ as well as the two M(IP)_2_C molecules M(IP)_2_C^ligand^ and M(IP)_2_C^OL^ around the transmembrane helices TMH11a and TMH12 was further analyzed (Fig. 7).

**Figure 7.**
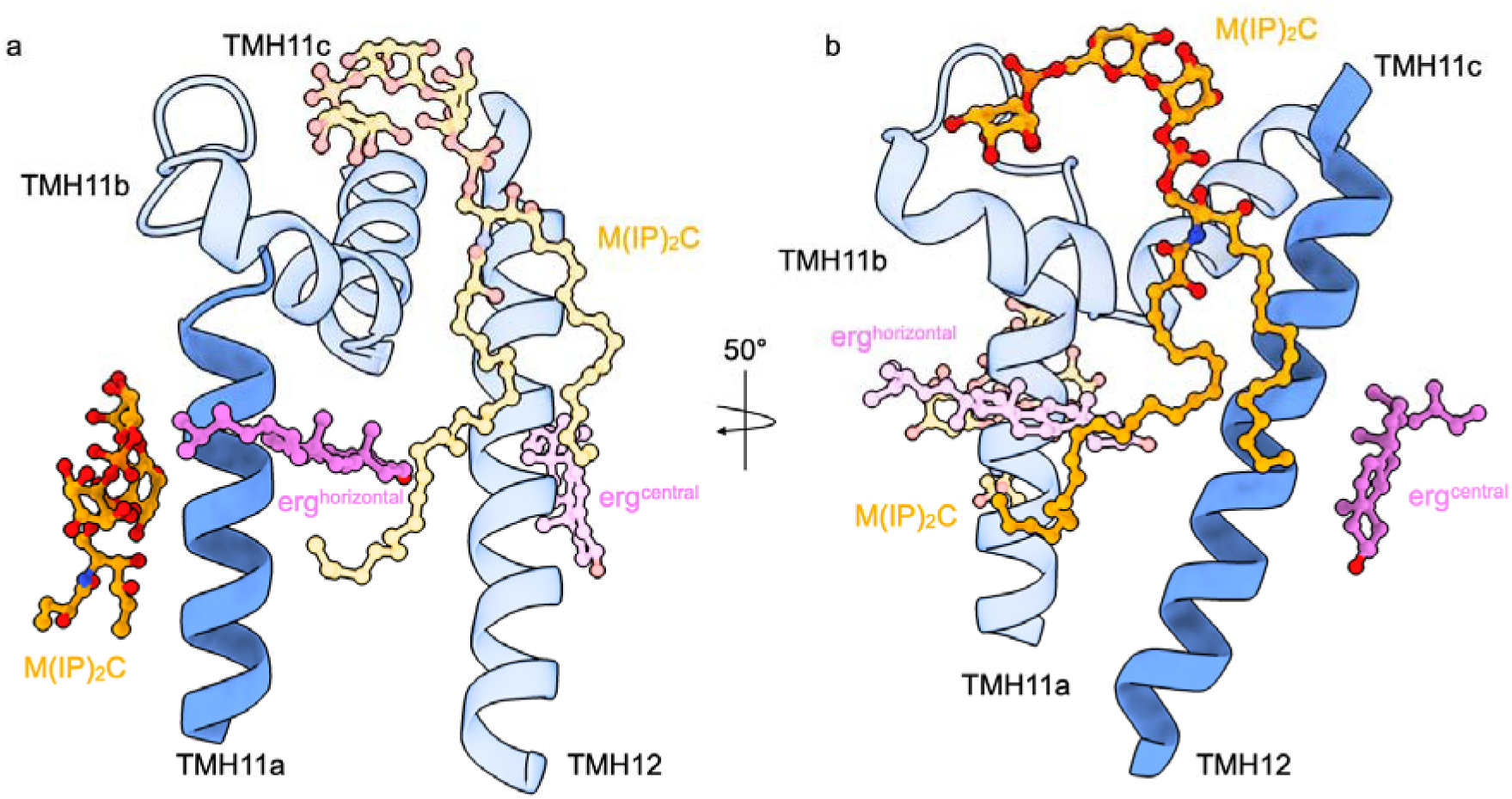
Pdr5-mediated interactions between the two M(IP)_2_C molecules and the two ergosterol molecules, erg^central^ and erg^horizontal^, respectively. (a) Focus on TMH11a connecting the M(IP)_2_C^ligand^ molecule in the substrate-binding site and erg^horizontal^. All transmembrane helices are shown as blue cartoons. TMH11a is displayed in full opacity, whereas TMH11b, TMH11c, and TMH12 are shown as transparent cartoons to help visualization. M(IP)_2_C and ergosterol molecules are shown in ball-and-stick representation. M(IP)_2_C^ligand^ located in the substrate-binding site is shown in orange, while M(IP)_2_C^OL^ located at the outer leaflet facing side of Pdr5 is shown in light orange. Erg^horizontal^ is shown in magenta, while erg^central^ is shown in light magenta. (b) Focus on TMH 12 connecting the M(IP)_2_C^OL^ molecule located at the outer leaflet facing site of Pdr5 and erg^central^. For ease of visualization, TMH12 is displayed in full opacity, whereas TMH11 is shown as transparent cartoon. M(IP)_2_C^OL^ located at the outer leaflet facing site of Pdr5 is shown in orange, M(IP)_2_C^ligand^ located in the substrate-binding site is shown in light orange. Erg^central^ is shown in magenta, while erg^horizontal^ is shown in light magenta.

Interestingly, TMH11 is split into three parts, a feature also observed in other members of the ABCG transporter family ^24, 39^. Consequently, TMH11a and TMH11c are in a perpendicular orientation to the membrane in Pdr5, while TMH11b, the middle part of this α-helix, makes an almost 90° bend inside the membrane.

M(IP)_2_C^ligand^ located in the substrate-binding site of Pdr5 mainly interacts with TMH11a. Erg^horizontal^ is positioned right at the opposite side of this helix and contacts both TMH11b and TMH12. TMH12 simultaneously engages with erg^central^ and the M(IP)_2_C^OL^ on the TMD-surface of Pdr5. In addition to the interaction with TMH12, M(IP)_2_C^OL^ also interacts with residues in TMH11a–c as well as the erg^horizontal^.

This formation of an extended sphingolipid-ergosterol interaction network connects M(IP)_2_C and ergosterol binding through the protein and suggests a possible molecular cross-talk coordinating binding of both, ergosterol and M(IP)_2_C.

## Discussion

The ABC transporter Pdr5 from *S. cerevisiae* presents a valuable model to investigate multidrug resistance mechanisms in pathogenic fungi. Here we report the cryo-EM structure of Pdr5 in complex with Mg^2+^/ADP and ATP at an improved resolution of 2.2 Å. By this, we were able to resolve additional densities that could be readily assigned as common constituents of the plasma membrane of *S. cerevisiae*, among which are four molecules of ergosterol ^30^. Numerous ergosterol molecules were also identified in a cryo-EM structure of the Pdr5 homolog Cdr1 from *C. albicans*, however, it was unclear which, and if, these molecules specifically interact with the transporter ^11^. We focused here on four ergosterol molecules that are located in distinct grooves that likely form stable interactions with Pdr5: erg^horizontal^ located horizontally in close proximity of the substrate-binding entry site, erg^central^ adopting a tilted conformation with respect to the membrane plane, and two ergosterols located in the outer leaflet, erg^OL1^ and erg^OL2^. Comparison of these ergosterol locations with the positioning of cholesterol around the human multidrug ABC transporter ABCG2, reveals a noteworthy similarity, though with slight changes ^31^. The cholesterol counterpart in ABCG2 to erg^horizontal^ is oriented vertically rather than horizontally (Fig. S7b) and is positioned at the level of the inner leaflet instead of the bilayer midplane. The equivalent cholesterol molecule in ABCG2 to erg^central^ is slightly shifted downwards (Fig. S7c) and likewise towards the inner leaflet. Erg^OL1^ occupies the same position as the cholesterol molecule in ABCG2. Finally, erg^OL2^ lies within the outer leaflet compared to the ABCG2 cholesterol in the inner leaflet, although arguably at the same height.

Despite these minor differences, both drug efflux pumps require the presence of a sterol for activity. For ABCG2, it was demonstrated that cholesterol acts as a positive regulator of the transporter, and depletion of membrane cholesterol reversibly reduces ABCG2 activity in mammalian cells ^40^. Likewise, we could show that the detergent DDM, when used for Pdr5 purification, dislodges the ergosterol molecules, ultimately resulting in a loss of ATPase activity (Fig. S8a). The failure of DDM to retain this sterol might explain why many previous biochemical characterizations of Pdr5 reported inactive protein when standard DDM-based protocols were employed ^34, 35^. For Cdr1 from *Candida albicans*, no sterols were detected following solubilization in DDM and subsequent detergent exchange to GDN^14^. In contrast, sterols were detected in Cdr1 from *Candida glabrata* preparations using PCC-α-M and dicarboxylate oside 9b (DCOD 9b)^11^, whereas sterol-like densities were observed in preparations of *Candida albicans* Cdr1 in the presence of LMNG and supplementation of cholesteryl hemisuccinate (CHS)^15^. Likely, only detergents that preserve some form of native ergosterol association yield functionally active Pdr5, which might be coupled to maintaining its overall structure within the detergent solubilized state.

To further address the specific role of ergosterol in Pdr5 functioning, its activity was tested in the presence of various amounts of ergosterol, revealing a concentration

dependent effect. Up until 15 % of ergosterol, Pdr5 activity proportionally increases with the ergosterol content. However, a further increase results in decreased activity, suggesting a modulation. This Gaussian-shaped increase then decrease in activity could be consistent with varying occupancy of multiple ergosterol-binding sites with distinct effects on protein activity. A similar principle has been described for the kinase GCN2, where partial inhibitor binding promotes kinase activity through positive allosteric coupling, whereas higher inhibitor concentrations result in inhibition ^41^. Such a dual regulatory effect could operate in Pdr5, explaining the different activities for ergosterol and cholesterol. As this modulating effect was not observed in the presence of cholesterol, it is ergosterol specific. Pdr5 shows a higher activity and catalytic efficiency with cholesterol than with its native sterol, ergosterol. For the human ABCG5/G8 transporter, ergosterol stimulated ATPase activity stronger than its physiological sterol, cholesterol ^42^. An effect that is also observed for Pdr5 but different to human ABCG2, for which cholesterol was reported to be the more potent activator ^23^. Structural and functional analyses of ABCG5/G8 identified a sterol-binding cavity and Y432 in ABCG5 involved in this differential response, suggesting that distinct sterols can differentially modulate ABC transporter activity ^42^. Y432 in ABCG5 occupies a different position as Y1478 in Pdr5 investigated in this study. Although these findings provide an example for higher ATPase activity by a non-native sterol, the physiological relevance of cholesterol-dependent effects on Pdr5 is limited by the fact that ergosterol, rather than cholesterol, is the predominant sterol in yeast membranes. Alternatively, the modulation of ergosterol might couple the ATP hydrolysis velocity and the induced conformational changes of Pdr5, *i.e*. inward- to outward-facing and back, to a most efficient substrate translocation. Furthermore, experiments performed with liposomes containing a mixture of both sterols revealed a dominance of ergosterol over cholesterol (Fig. 5). At saturating levels of ergosterol, further addition of cholesterol did not change the activity. In contrast, at a fixed concentration of cholesterol, addition of ergosterol modulated the activity towards saturating levels of ergosterol. Surprisingly, Pdr5 activity in presence of only cholesterol is elevated, but the higher activity cannot be sustained upon addition of ergosterol, thereby confirming the modulating effect of ergosterol. Furthermore, this indicates that Pdr5 binds ergosterol with higher affinity than cholesterol. To assess whether a single ergosterol, or overall ergosterol presence is responsible for the modulation of the activity, mutational studies were performed. By mutating the amino acid residues L1329 and Y1478, the erg^central^ binding-site was widened to enable better accessibility for cholesterol. This engineering approach induced modulation of Pdr5 activity in the presence of cholesterol, thereby pinpointing erg^central^ as modulation site. Interestingly, the conservation of 78 % for L1329 and 32 % for Y1478 (500 sequences of the PDR-subfamily aligned) suggests that, rather than the precise nature of the amino acid residues, shape complementarity and spatial constraints are the major determinants of this ergosterol binding site.

Placing the physiological role of this modulation into a biological context is challenging. There are multiple reports about the cellular content of ergosterol in *S. cerevisiae*. The amount of ergosterol, analyzed via MS, is described to be approximately 12 mol% ^30, 43^. However, this value represents the total ergosterol content of a yeast cell, not discriminating between the different membrane types in the organism. Additionally, cellular membranes are not symmetrical ^44^. Approximately 70 % of the cellular ergosterol pool in yeast is located in the plasma membrane, while the remaining 30 % reside in the endoplasmic reticulum. Within the plasma membrane, ergosterol is asymmetrically distributed across the bilayer, in an outer: inner leaflet ratio of ∼ 15:55, which means that roughly 80 % resides in the cytoplasmic leaflet, where it accounts for about 45 % of all the lipids in the inner leaflet ^32, 45^. Given the fact that Pdr5 appears to reside inside lipid microdomains ^19^, likely enriched in ergosterol ^46, 47^, the local ergosterol concentration might be even higher. Xenobiotic efflux by Pdr5 is an ATP-consuming process that comes at an energetic cost. It has been proposed that such basal activity keeps multidrug exporters in a state that allows rapid removal of potentially harmful compounds ^37, 48^. The modulation of Pdr5 activity by ergosterol may therefore link the activity of this energy-consuming efflux system to the state of the membrane. Reduced Pdr5 activity could decrease the energetic investment in xenobiotic efflux, while reducing the capacity for rapid removal of potentially harmful compounds. Such coupling could be relevant for balancing the energetic cost of maintaining xenobiotic efflux capacity with the need for rapid protection against potentially harmful compounds.

Going one step further, previous studies demonstrated that Pdr5 can mediate the transport of steroid molecules such as estradiol ^49^. This led to the hypothesis, that endogenous sterols like ergosterol might be a physiological substrate of Pdr5. However, these studies did not directly prove ergosterol to be transported by Pdr5 and the hypothesis was a derivation from the experiments performed with estradiol. Our results are consistent with an alternative model in which ergosterol modulates the activity of Pdr5 through protein-lipid interactions rather than acting as a substrate of the transporter.

In addition to the localization, the composition of the lipid bilayer is functionally relevant as well. In *S. cerevisiae*, Pdr5 was shown to confer reduced resistance in ergosterol-deficient backgrounds which could not be explained by enhanced membrane permeability nor changed fluidity ^27^. Previous work performed with heterologously expressed Cdr1 in *S. cerevisiae* showed mis-localization of the protein in backgrounds with deletions in the ergosterol biosynthetic pathway leading to reduced functionality ^50^. A more recent study investigating the effect of ergosterol deficiency in *C. albicans* on Cdr1 and Pma1 demonstrated a more differentiated result ^51^. These differences may be related to differences in the metabolism of these two organisms, although *S. cerevisiae* serves as a suitable model organism. Reduced levels of ergosterol altered the association of Cdr1 with the plasma membrane and membrane microdomains and were associated with partial delocalization and increased drug susceptibilities. The existence or structural integrity of lipid microdomains was not necessarily crucial to retain Cdr1 in the plasma membrane. Another study investigated the effect of altered ergosterol content on the functionality of the yeast efflux pump Snq2 and Pdr5 ^18^. While the activity of Snq2 was almost unaffected by altered ergosterol levels/ composition, the activity of Pdr5 was highly impaired. The protein remained properly localized to the plasma membrane and Pdr5 overexpression did not substantially rescue the phenotype. This supports our observation that ergosterol modulates Pdr5 activity through mechanisms beyond general changes in membrane properties.

Besides ergosterol, also two molecules of the sphingolipid M(IP)_2_C could be identified for the first time in the cryo-EM structure of an ABC transporter. Especially, the M(IP)_2_C^ligand^ molecule located at substrate-binding site of Pdr5 catches the eye, as the localization of the M(IP)_2_C headgroup shows remarkable resemblance with the well-studied Pdr5 substrate R6G (Fig. S6). M(IP)_2_C is synthetized in the Golgi and subsequently transported to the plasma membrane via the vesicle-mediated secretory pathway, as no dedicated transporter for M(IP)_2_C has been reported to date ^52, 53^. The substrate-binding site of Pdr5 is predominantly hydrophobic and extends into the membrane, with access from the inner leaflet ^10^. Interestingly, the detergent LMNG was observed in the substrate-binding cavity of Cdr1 from *Candida albicans* ^13^. This observation, together with the mainly hydrophobic nature of the substrates of Pdr5 ^10^ indicates that lipids could occupy this cavity. In addition, Pdr5 has been linked to plasma membrane lipid organization through genetic interactions with the phospholipid flippases Dnf1/Lem3 and Dnf2/Lem3 ^54^. Whether these observations are related and whether lipids such as M(IP)_2_C can directly interact with the Pdr5 substrate-binding cavity remains unclear. Further investigations will be required to elucidate our observation.

The presence of sphingolipids is coherent with our observation that Pdr5 localizes into DRMs. The Pdr5 ortholog Cdr1 from *Candida albicans* was shown to not only be located in lipid microdomains enriched in ergosterol and sphingolipids but its targeting into these microdomains as well as its activity depends on these membrane constituents^47, 51^. The requirement for ergosterol and sphingolipids might represent a broader feature among fungal ABC transporters. However, whether similar sphingolipid dependencies contribute to Pdr5 localization and activity, remain to be investigated. In that respect, the arrangement of erg^horizontal^, erg^central^ and the two M(IP)_2_C molecules around TMH11 and TMH12 (Fig. 7) identifies these helices as a center for lipid engagement in Pdr5. The finding that these two helices simultaneously contact the substrate-binding site erg^horizontal^, the peripheral erg^central^ and the two M(IP)_2_C molecules suggests that TMH11 and TMH12 could serve as an interface linking the surrounding lipid environment to the substrate-binding region of Pdr5.

Within this extended network, four residues of Pdr5 have been functionally characterized previously (Fig. S13). Residue S1360 lines the substrate-binding cavity and was identified as a determinant of substrate specificity and R6G binding ^55^ and confirmed as a binding-site residue in the Pdr5 structure ^10^. Residue T1364 was recovered among FK506-resistance suppressor mutations as part of the T1364S/ M1379V variant ^56^. Amino acid M1365 is oriented away from the substrate-binding cavity and towards the side chain of erg^horizontal^ and tolerates substitutions without a distinct phenotype ^57^. W1383, described as a FK506-resistant suppressor mutant as well ^56^, contacts the erg^horizontal^ lipid tail together with amino acid residue M1365. Although this amino acid residue is highly conserved (99.6 % among the PDR family), this conservation more likely reflects its position at the interface between the two halves of the transporter. The remaining amino acid residues in the network, although partially highly conserved, have not been characterized previously. While this structural connection could provide a basis through which Pdr5 senses or responds to its lipid environment, further studies will be necessary to validate this proposed mechanism.

Taken together, our results highlight the importance of the membrane environment for Pdr5 function and demonstrate that ergosterol is a specific and functionally relevant component of the Pdr5 membrane environment. We show that ergosterol binds to distinct grooves in the protein, modulates its activity and that cholesterol cannot substitute for ergosterol to maintain Pdr5 activity. Our findings provide new insights into the role of membrane lipids in the regulation of this ABC transporter.

## Material & Methods

### Chemicals

All chemicals, if not stated otherwise, were obtained from Sigma-Aldrich, Carl Roth, VWR or Thermo Fisher Scientific. The phospholipids POPE, POPC and POPS were purchased from Avanti Polar Lipids. Ergosterol and PCC-α-M were purchased from Fluka and Glycon Biochemicals, respectively. Bio-Beads SM-2 beads were purchased from Bio-Rad Laboratories and prepared as described by Geertsma et al58.

### Yeast Strains and Plasmid Mutagenesis

The *Saccharomyces cerevisiae* strains used in this study are listed in Table 1 and were generated via site-directed mutagenesis of *pdr5* on plasmid pRE5 ^59^. Details about plasmid and strain construction can be found in Table 2.

**Table 1.** List of yeast strains used in this study.

| Strain | Genotype |
| --- | --- |
| YALF A1 (parent strain, Pdr5 WT) | MATa; *ura3-52*; *trp1-1*; *leu2-3,112*; *his3-11,15*; *ade2-1*; *pdr1-3*<br>MATa; *ura3-52*; *trp1-1*; *leu2-3,112*; *his3-11,15*; *ade2-1*; *pdr1-3*; ΔPDR5::TRP1 |
| YRE1001 (ΔPP) | MATa; *ura3-52*; *trp1-1*; *leu2-3,112*; *his3-11,15*; *ade2-1*; *pdr1-3*; ΔPDR5, ΔPDR5prom::TRP1 |
| 14xHis-Xa-Pdr5-N734A | MATa; *ura3-52*; *trp1-1*; *leu2-3,112*; *his3-11,15*; *ade2-1*; *pdr1-3* |
| 14xHis-Xa-Pdr5-L1329A-Y1478A | MATa; *ura3-52*; *trp1-1*; *leu2-3,112*; *his3-11,15*; *ade2-1*; *pdr1-3* |
| 14xHis-Xa-Pdr5-E1036Q | MATa; *ura3-52*; *trp1-1*; *leu2-3,112*; *his3-11,15*; *ade2-1*; *pdr1-3* |

**Table 2.**
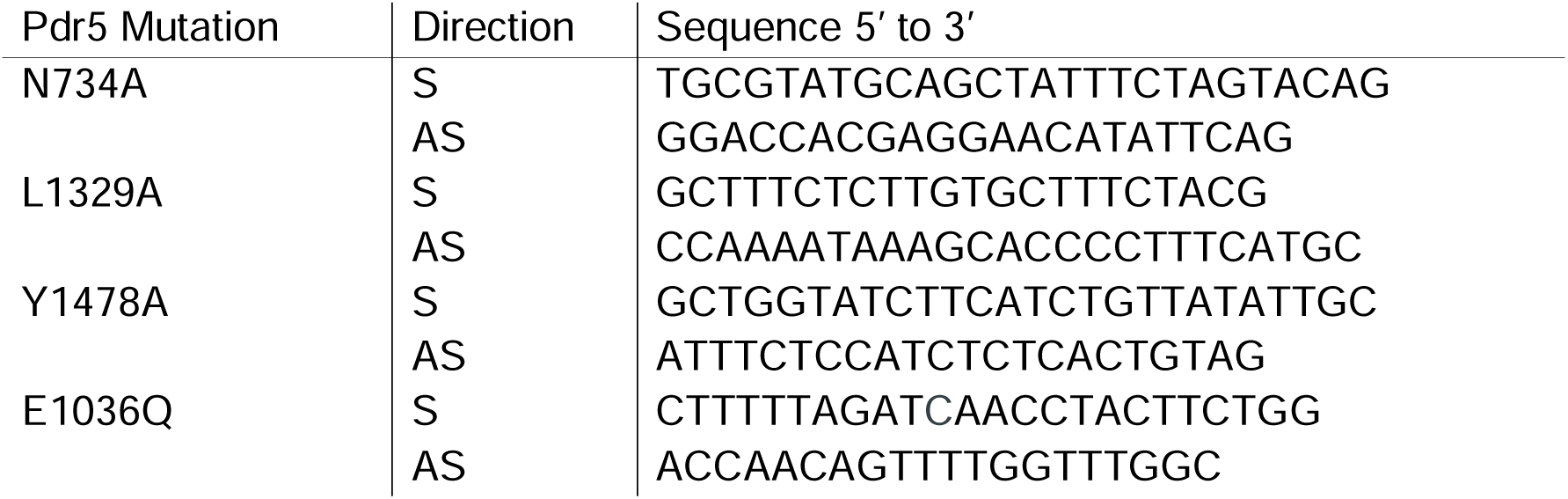
List of primers used for introducing point mutations in *pdr5* gene on pRE5 plasmid. *S is sense, and AS is antisense*.

### Protein expression and purification

Pdr5 and the desired mutations were expressed and purified as described in Wagner et al. ^34^ In short, the *S. cerevisiae* strain expressing the desired Pdr5 variant was incubated in YPD medium (20 g/l tryptone/ peptone, 10 g/l yeast extract, 20 g/l glucose) at 25 °C and 140 rpm. At an OD_600_ of 1.5, the nitrogen source was replenished by the addition of a 10th volume of 5x YP medium (50 g/liter yeast extract; 100 g/liter tryptone/peptone). Cells were harvested at an OD_600_ of 3.5 to 4.0. The cells were disrupted with glass beads and cell debris removed via differential centrifugation at 4 °C (2x 1,000 xg for 5 min, followed by 3,000 xg for 10 min). The membranes were pelleted at 20,000 xg and 4 °C for 40 min and resuspended in buffer A (50 mM Tris-HCl, pH 7.8, 50 mM NaCl, 10 % (w/v) glycerol). The membranes, adjusted to a total protein concentration of 10 mg/ml, were solubilized under gentle stirring with 1 % (w/v) PCC-α-M at 4 °C. The solubilized protein was separated from the non-solubilized material by ultracentrifugation at 170,000 xg and 4 °C for 1 h. The detergent concentration of the supernatant was adjusted to a concentration of 0.55-0.60 % (w/v) prior to loading onto a 1 ml HiTrap Chelating HP column, loaded with Zn^2+^ ions. The column was pre-equilibrated with low histidine buffer (50 mM Tris-HCl, pH 7.8, 500 mM NaCl, 10 % (w/v) glycerol, 2.5 mM L-histidine, 0.003 % (w/v) PCC-α-M). The protein was washed and eluted using a step gradient with low and high histidine buffer (50 mM Tris-HCl, pH 7.8, 500 mM NaCl, 10 % (w/v) glycerol, 100 mM L-histidine, 0.003 % (w/v) PCC-α-M).

For size exclusion chromatography, elution fractions were pooled and concentrated with a Vivaspin^TM^ Turbo 15 (100 kDa MWCO). The concentrated protein sample was further purified on a Superdex 200 Increase 10/300 GL column equilibrated with buffer A containing 0.003% (w/v) PCC-α-M.

For the quantification of ergosterol molecules in the Pdr5 sample purified in DDM, the protein was treated as described above. The membranes were solubilized in 1 % (w/v) DDM under gentle stirring at 4 °C. Buffers were supplemented with 0.05 % DDM. Elution fractions after affinity purification were pooled, concentrated and used for lipid extraction and MS.

### Ergosterol quantification in the purified protein sample

For the quantification of the number of ergosterol molecules per Pdr5 molecule, 0.03 mg (0.18 nmol) purified Pdr5 in detergent were adjusted to final volume of 100 µl. Prior to GC–MS analysis, 0.5 nmol of internal standard β-sitosterol was added. Lipids were extracted with 0.3 ml n-butanol (2 times) and solvent evaporated with nitrogen gas. Dried lipid films were resuspended in 100µl MSTFA and incubated (80 °C; 30 min). 1 μl of derivatized compounds was injected and measured on a 5977B GC/MSD (Agilent Technologies) as described ^60^. Oven temperature gradient: constant at 70 °C for 1 min, ramped at 42 °C min^−1^ to 280 °C, further with 4 °C min^−1^ to 320 °C, held constant for 3 min (total 19 min). Electron ionization source: 70 eV; source temperature: 200 °C, mass range: 60-600 *m*/*z*, 5 scans per second. Sterols were identified *via* MassHunter Qualitative (v b08.00, Agilent Technologies) by NIST14 Mass Spectral Library comparison (https://www.nist.gov/srd/nist-standard-reference-database-1a-v14). Peaks were integrated using MassHunter Quantitative (v b08.00, Agilent Technologies). The sterol content was quantified by using a calibration with β-sitosterol as internal standard (Merck Sigma-Aldrich). The quantitative amounts were further normalized to the Pdr5 content.

### Cryo-EM sample preparation and data collection

Pdr5 reconstituted in peptidiscs (1.5 mg/mL) was mixed with 2 mM of ATP and 2 mM MgCl_2_ and incubated for 2 min. The solution was mixed by pipetting and used immediately for grid preparation. 3 µL of the sample mixture was applied onto an Au 300-mesh EM grid with R1.2/1.3 UltrAuFoil support film (Quantifoil Micro Tools GmbH, Großlöbichau, Germany) which had been glow-discharged prior to use. The sample applied to the grid was vitrified in liquid ethane at –180℃ using a Vitrobot Mark IV (Thermo Fischer Scientific Corp., Eindhoven, The Netherlands).

The grids were imaged using Titan Krios G3 TEM (Thermo Fischer Scientific Corp., Eindhoven, The Netherlands) operated at liquid nitrogen temperature, 300-kV accelerating voltage, in EF-TEM mode with GIF slit size of 20 µm, and housed at eBIC facility (Titan Krios I, M02, Diamond Light Source Ltd, Didcot, Oxfordshire, UK). The micrographs were recorded using automated image acquisition software EPU (Thermo Fischer Scientific Corp., Eindhoven, The Netherlands) on a K3 direct electron detector (Gatan Inc., Pleasanton, CA, USA) in super-resolution bin-2 mode, at a detector pixel size of 0.831 Å (nominal magnification 105,000×) and nominal defocus values between –0.7 and –2.2 µm. The exposures lasted 1.9 s, collected in 70 fractions and fluency of 48.9 e^−^/Å^2^ or 17.77 e^−^/px/s. This dataset contained micrographs.

### Cryo-EM data processing for structure solution

The cryo-EM map of Pdr5 was reconstructed in cryoSPARC v4.7 ^61^ (Table S1). Micrographs were corrected for beam-induced sample motion and contrast transfer function (CTF) parameters were estimated on patched micrographs. A subset of the images with estimated maximum resolution below 4 Å was selected from the full dataset for further processing. 2,000 manually picked particles were used to prepare a set of 2D references for template-based particle search, which was subsequently optimized to yield diverse projections of Pdr5. The initial particle pool was extracted in 380-pixel boxes and pruned through a series of reference-free 2D classification rounds to 632,196 particles. A model for cryo-EM reconstruction was created through *ab initio* volume reconstruction algorithm. The model and the pruned particle pool were subjected to 3D auto-refinement, which generated the first intermediate cryo-EM map used for subsequent 3D classification and refinement. The final map reconstruction was performed using non-uniform refinement (NU-refinement) algorithm in cryoSPARC ^62^. The overall resolution of the obtained map, as judged by Fourier-shell correlation (FSC) of the half-maps with the gold-standard FSC_0.143_ criterion ^63^, was 2.45 Å.

### Model building and refinement

The IF conformation of Pdr5 bound to ATP and ADP (pdb entry 7P04) ^10^ was docked into the reconstructed cryo-EM map using ChimeraX ^29^. The obtained model was adjusted manually in Coot-0.9 ^64^. Model refinement was carried out in real space using PHENIX-2.0 ^65^. To assist model building, the maps were subjected to a density modification procedure ^65^, which improved resolution to 2.2 Å. Ligands were manually docked into the density after the protein model was finally placed in the cryo-EM map. Locally sharpened maps were also used for visualization purposes. Molecular graphics were rendered in ChimeraX-1.10 ^29^. All refinement and data collection statistics are summarized in Table S1.

### Native MS and top-down MS analysis

In order to perform the native MS experiments, Pdr5 Asn^734^Ala was generated in which the only glycosylation site in Pdr5 had been removed. This protein was highly phosphorylated and was subjected to λ-phosphatase treatment. Pdr5 WT was used for cross-verification.

The chromatogram resulting from the native top-down experiments identified several lipid species in the range of 1300-1400 Da pointing towards M(IP)_2_C, an abundant sphingolipid in yeast ^30^. Several species with different lengths and hydroxylation of the ceramide chain are present in this chromatogram (M(IP)_2_C 44:0;O2, M(IP)_2_C 44:0;O3, M(IP)_2_C 44:0;O4, M(IP)_2_C 46:0;O3, and M(IP)_2_C 46:0;O4). Among these M(IP)_2_C 44:0;O4 was the most dominant species with a molecular weight of 1359 Da (Fig. S5). After ergosterol, M(IP)_2_C 44:0;O4 is the most abundant lipid species in yeast ^30^.

Both wild type PDR5 and Asn^734^Ala were treated with *λ*-phosphatase (New England Biolabs) to reduce the phosphorylation before buffer-exchanged into 500 mM ammonium acetate, pH 7.5, containing 0.1% (w/v) Octyl Glucose Neopentyl Glycol (Anatrace) using a Zeba Spin 40K MWCO desalting column (Thermo Scientific). Protein concentration was determined using NanoDrop One and adjusted to the desired concentration for native mass spectrometry analysis. Typically, 3 μL of desalted protein was loaded into an in-house prepared gold-coated capillary and electrosprayed into a Q Exactive UHMR hybrid quadrupole-Orbitrap mass spectrometer (Thermo Fisher Scientific) using the following settings: capillary voltage, 1.4 kV; UHV pressure, 7.5 × 10^−10^ mbar; capillary temperature, 200 °C; and resolution, 17,500. Unless otherwise stated, all experiments were performed in positive mode. Raw MS data were processed with Xcalibur 4.1.50 (Thermo Fisher Scientific), and spectra were deconvoluted using UniDec v6.0.1 ^66^.

To identify the co-purified lipids, we performed the native top-down experiments as previously reported ^67,68^. Typically, desalted protein was loaded into a gold-coated capillary and electrosprayed into an Orbitrap Eclipse Tribrid mass spectrometer (Thermo Fisher Scientific). MS^1^ spectra were collected at the resolution of 15,000 at *m/z* = 200. For the pseudo-MS^2^ experiments, protein ions were isolated in the ion-trap (50 *m/z* isolation window with a *Q* value of 0.125) and lipids were released with high in source collision energy (SID = 250, and HCD = 200). For the pseudo-MS^3^ experiments, released precursor ion was isolated in quadrupole using a 2 *m/z* window, and subjected to HCD (15 V absolute), followed by *m/z* analysis in the Orbitrap at a resolution of 240,000 at *m/z* = 200.

### Isolation of detergent resistant membranes

Detergent resistant membranes were isolated as described by Malínská et al. ^69^ with modifications. The *S. cerevisiae* strains expressing the desired Pdr5 variant were used to inoculate a preculture in YPD medium and were incubated at 30 °C and 180 rpm overnight. Subsequently, a main culture was inoculated to an OD_600_ of 0.1 and grown to a final OD_600_ of 0.6-0.8. An equivalent to 50 OD_600_ units were harvested, washed with ice-cold MQ and the cell pellets resuspended in 1 ml TNE buffer (50 mM Tris-HCl, pH 7.8, 150 mM NaCl, 5 mM EDTA) supplemented with 4 µM leupeptin and 2 µM pepstatin. Cells were disrupted with glass beads (Ø 0.5 mm) on a Disruptor Genie^TM^ twice for 5 minutes at 4 °C, with a 5-minute incubation on ice between cycles. The lysate was separated from the glass beads and unbroken cells and cell debris removed via centrifugation. The supernatant was centrifuged to collect the crude membranes. Membranes equivalent to 50 OD_600_ units were resuspended in 225 µl TNE buffer. Triton X-100 was added to a final concentration of 1 % and the samples incubated on ice for 30 minutes. Following incubation with Triton X-100 on ice, 500 µl Optiprep (STEMCELL^TM^ Technologies) were added to the samples to a final concentration of 40 % (w/v) and transferred to a centrifuge tube. The mixture was overlaid sequentially with 1 ml of 30 % (w/v) Optiprep in TXNE buffer (TNE buffer supplemented with 0.1 % Triton X-100), 1 ml of 20 % (w/v) Optiprep in TXNE buffer, 250 µl of TXNE buffer, followed by centrifugation at 248,000 xg and 4 °C for 2 hours.

Using a peristaltic pump, six equal fractions were collected from the bottom of each gradient and proteins precipitated with 15 % (v/v) trichloroacetic acid. Protein pellets were washed with ice-cold acetone, dried and resuspended in SDS sample buffer. The samples were analyzed via SDS-PAGE, immunoblotting and LC-MS.

### SDS-PAGE and immunoblotting

For analysis of Pdr5 samples by SDS-PAGE and Western Blot, 5-10 µg protein solubilized in SDS-PAGE sample buffer were separated by electrophoresis before transferring to a PVDF membrane (0.45 µm, Amersham Hybond) at constant current. The membranes were blocked overnight in blocking solution (10 % milk powder, 0.05 % sodium azide (Sigma) in TBS, 0.1 % Tween 20 (TBS-T)). Blocking solution was removed by washing with TBS-T. The Pdr5-specific primary antibody ^70^ was diluted 1:3000 in TBS-T substituted with 3 % BSA. The blots were incubated in the primary antibody for 1 hour at room temperature under gentle agitation. The blots were washed with TBS-T before addition of the secondary antibody. The secondary antibody (goat, Anti-Rabbit IgG (whole molecule)–peroxidase antibody, Sigma) was diluted 1:10000 in TBS-T containing 5 % milk powder. The blots were incubated at room temperature for 1 hour under gentle agitation and subsequently washed with TBS-T and TBS. The blots were developed with a WESTAR ηC Ultra 2.0 chemiluminescent substrate kit (Cyanagen) and the signals detected with an Amersham Imager 680.

### LC-MS analysis of detergent resistant membranes

50 µl of each fraction (see isolation of detergent resistant membranes) were spiked with 10 µg of the internal standard n-Dodecyl β-maltoside (DDM), extracted twice with 300 µl of n-butanol, dried and resuspended in 50 µl of methanol. Samples were analyzed using an Agilent Technologies 1290 Infinity high-performance liquid chromatography (HPLC) system, coupled to a heated electrospray ionization–mass spectrometer (Thermo Q Exactive Plus; Thermo Fisher Scientific). 5 µl was injected into an ACQUITY™ UPLC CSH C18 1.7 µm Column, 2.1 × 150 mm (Waters Chromatography Ireland Ltd) operating at 55 °C with a flow rate of 300 µl/min. Separation of the compounds was achieved by a changing gradient of eluent A (5 mM ammonium formate in water/acetonitrile 40:60, v/v) and eluent B (5 mM ammonium formate in acetonitrile/2-propanol, 10:90, v/v). The following linear gradient was applied: 1) 45% eluent B for 2.5 min, 2) a gradient from 45% to 90% eluent B over 19.5 min, 3) holding for 3 min, 4) returning to 45% eluent B in 0.5 min, 5) and holding for 8 min. The column effluent was injected directly into the Thermo Q Exactive Plus operating in negative ion mode: Spray Voltage: 3.20 |kV|, Capillary temperature: 230 °C, S-lens RF level: 50.0, Sheath gas flow: 30, Auxiliary gas flow: 20, Sweep gas flow: 3, Aux gas heater temperature: 380 °C. Spectral data constituting total ion counts were analyzed using the the Thermo Scientific XCalibur processing software by applying the Genesis algorithm-based automated peak area detection and integration. The total ion counts of the extracted lipid products: IPC (*m/z* 952.6860 [M-H]^-^) and MIPC (*m/z* 1114.7388 [M-H]^-^) were corrected for the internal standard DDM (*m/z* 509.2962 [M-H]^-^) and plotted as normalized ion count on the Y-axis.

### Preparation of liposomes

Liposomes were prepared and Pdr5 reconstituted as described by Gala Marti et al ^37^. Stock solutions of POPC, POPE, POPS and sterol in chloroform were mixed in the desired ratio, using a total lipid mass of 40 mg per sample. Lipid compositions are reported as weight percentages (% (w/w)) relative to the total lipid mass. Sterol content was varied between 5 and 20 % (w/w), while the remaining lipid fraction consisted of POPC, POPE, and POPS in fixed relative mass ratios (30:30:20), adjusted proportionally to accommodate changes in sterol content.

To compare the effects of ergosterol and cholesterol, liposomes were prepared as described above with either sterol. Unless stated otherwise, “sterol” refers to ergosterol or cholesterol used interchangeably at identical weight fractions.

Using a rotary evaporator, chloroform was removed at 40 °C and 150 rpm for 60 min. The lipid film was suspended in buffer A without glycerol (50 mM Tris-HCl, pH 7.8, 50 mM NaCl) to a homogenous suspension and adjusted to a final lipid concentration of 20 mg/ml. The suspension was subjected to six cycles of sonication (5 s on/ 5 s off) in an ice-water bath to obtain small unilamellar vesicles (SUVs). To generate large multilamellar (LMVs), the SUV-suspension was flash-frozen in liquid nitrogen and thawed at room temperature for five times. The resulting LMVs were stored at -80 °C.

### Reconstitution of Pdr5 in liposomes

To generate large unilamellar vesicles (LUVs), the LMVs were thawed at room temperature and extruded 11 times through a 400 nm polycarbonate filter. The LUV-suspension was diluted to a final concentration of 4 mg/ml with buffer A without glycerol. To incorporate purified Pdr5 into the liposomes, the liposomes were destabilized by the addition of small volumes of 10 % (w/v) Triton X-100 while monitoring the absorption at 540 nm. Addition of detergent was continued until the absorption reached ∼60 % of the initial value.

Purified Pdr5 was added to the liposomes in a protein-to-lipid ratio of 1:100 (w/w) and incubated at room temperature under gentle agitation. Bio-Beads SM-2 were added to obtain 200 mg per 5 ml suspension and incubated at 4 °C under gentle agitation for 20 min. The Bio-Beads were added twice more and the mixture was incubated overnight at 4 °C under gentle agitation. On the following day, an additional amount of Bio-Beads was added and the suspension incubated at 4 °C under gentle agitation for further 60 min prior to removal of the beads using a disposable Poly-Prep chromatography column.

### Determination of reconstitution efficiency and orientation of protein in liposomes

The efficiency of the protein reconstitution and the orientation of the ABC-transporter in the proteoliposomes was determined as described by Gala Marti et al. ^37^ using a quantitative SDS-PAGE. Preceding the analysis, the proteoliposomes were concentrated five-fold by ultracentrifugation at 220,000 xg and 4 °C for 20 min. The proteoliposomes pellet was resuspended in buffer A without glycerol.

To accommodate the samples with higher amounts of sterol, a modified SDS sample buffer (100 mM Tris-HCl, pH 6.8, 0.02 % (w/v) bromophenol blue, 10 % (w/v) SDS, 8 M Urea) was prepared as 3x stock and mixed with the samples in a 2:1 ratio (buffer: protein sample).

For the determination of the reconstitution efficiency, 30 µl of the proteoliposomes supplemented with the 3x SDS sample buffer were applied onto a 7 % polyacrylamide gel together with 10 µl of a BSA standard of known concentration and separated by electrophoresis. The SDS-PAGE gel was stained with Quick Coomassie Stain and the protein bands were quantified using ImageJ software ^71^.

The fraction of the protein reconstituted in an inside-out orientation was determined by a proteolytic digestion assay using trypsin. A volume of 40 µl of the five-fold concentrated proteoliposomes was incubated with 2 µl of 0.1 mg/ml trypsin stock in 1 mM hydrochloric acid for 4 h at 37 °C. As a control, 1 % (v/v) Triton X-100 was added prior to the trypsin treatment to solubilize the liposomes. The samples were supplemented with the 3x SDS sample buffer and subsequently analyzed by 7 % SDS-PAGE as described above.

### Measurements of the anisotropy and general polarization

To assess whether the substitution of ergosterol in the liposomes with cholesterol impacts the overall membrane characteristics, anisotropy, an indicator for membrane fluidity, and the general polarization, which reflects the degree of membrane packing, were determined.

The measurements were performed using Fluorolog III fluorescence spectrometer (Horiba) in a quartz glass cuvette. Extruded liposomes (LUVs) with varying ergosterol or cholesterol content (100 µM lipid content) were mixed with Laurdan (0.3 µM) to obtain a dye: lipid ratio of (1:333). The mixture was incubated at 37 °C and 300 rpm for 1 h in the dark. For the measurement, 1 ml of this sample was used.

For the anisotropy measurements, the emission was recorded at 435 nm (5 nm slit width) while exciting at 350 nm (5 nm slit width).

The anisotropy was calculated using Equation 1:

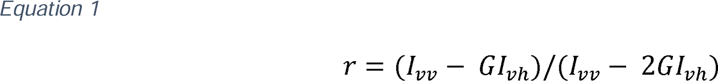

where is the measured fluorescence intensity, are the vertical and horizontal settings for the excitation and emission polarizers, respectively, and is the instrumental correction factor (provided by each instrument).

General polarization was determined by exciting the sample at 350 nm (3 nm slit width) and recording an emission spectrum from 400-600 nm (3 nm slit width).

The general polarization was calculated with Equation 2 using the fluorescence intensity at the indicated wavelength:

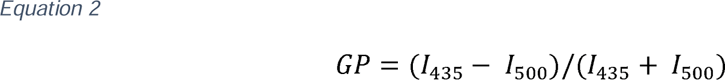

### ATPase activity

The ATPase activity of Pdr5 in liposomes containing various amounts of ergosterol or cholesterol was determined using a colorimetric assay with malachite green. The proteoliposomes were incubated in buffer A without glycerol substituted with or without 5 mM magnesium chloride and varying ATP concentrations (0-5 mM) in a final volume of 50 µl. After incubation at 30 °C for 60 min, the reaction was stopped by addition of 25 µl of the reaction mix to 175 µl of ice-cold 40 mM sulfuric acid ^72^. The amount of released inorganic phosphate was determined using Na_2_HPO_4_ as a standard ^73^.

The measured ATPase activity was corrected for the determined (i) efficiency of protein reconstitution into the liposomes and (ii) the fraction of ABC-transporter with an inside-out orientation in the proteoliposomes.

The ATPase activity of detergent-solubilized Pdr5 was determined as described by Wagner et al. ^34^.

## Acknowledgements

We thank all members of the Institute of Biochemistry, HHU, for stimulating discussions. We are also grateful for support of Cigdem Günes and Ursula Fleig. This work was supported by the Deutsche Forschungsgemeinschaft DFG (grant Schm1279/17-1 to L.S.). Furthermore, this work was supported by ERC Advanced Award (742210) and Wellcome Trust Investigator Award (222451/Z/21/Z) Advanced (to B.F.L.). The Center for Structural Studies is part of StrukturaLink Rhein-Ruhr which is funded by the Deutsche Forschungsgemeinschaft (DFG Grant number 573727698 to S.S. and INST 208/749 1 FUGG). Lipidomics analyses were conducted in the CEPLAS metabolism and metabolomics laboratory (CMML), which is supported by the Cluster of Excellence on Plant Sciences (CEPLAS) under Germanýs Excellence Strategy EXC-2048/1 under project ID 390686111.

## Data availability

Cryo-EM maps and coordinates have been deposited in the EMDB and PDB, respectively, with accession numbers EMD-100182 and PDB 34VA. The mass spectrometry raw data have been deposited in the Figshare database.

## Authors contributions

Conceptualization: C. R., B. F. L., S. H. J. S., L. S., Funding acquisition: L.S., Methodology: C. R., M. E., A. H., Investigations: L.-M. N., J. H., E. K., S. K., H. S., M. E. Writing original draft: L.-M. N., J. H., L. S., Writing review and editing: all authors.

