## Supplementary Information for "The multidrug transporter Pdr5 from *Saccharomyces cerevisiae* is an ergosterol-modulated efflux pump"

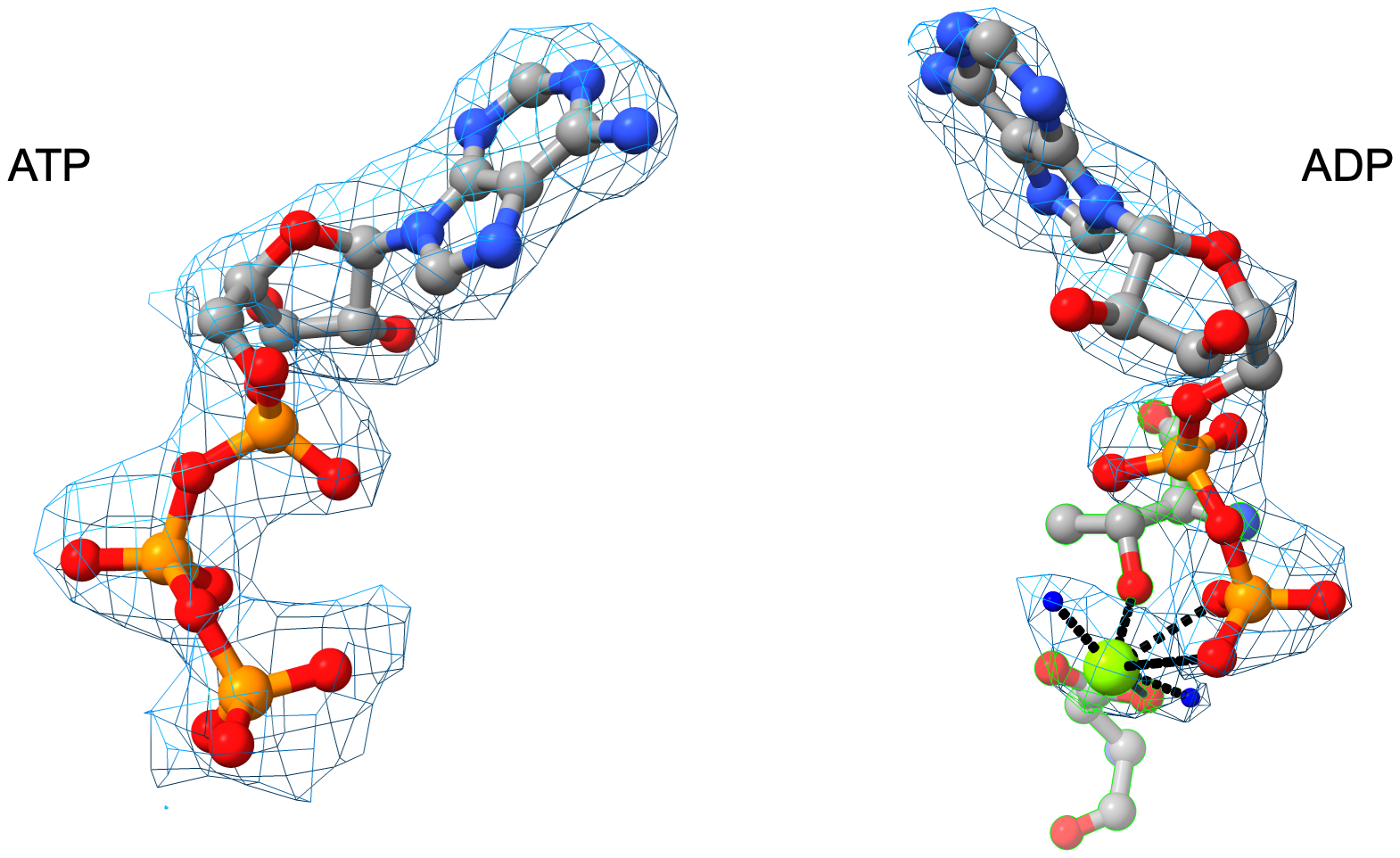

**Figure S1** Portions of the cryo-EM map showing the bound nucleotides in the Pdr5 structure, ATP in the deviant NBD1 and ADP/ Mg^2+^ in the canonical NBD2.

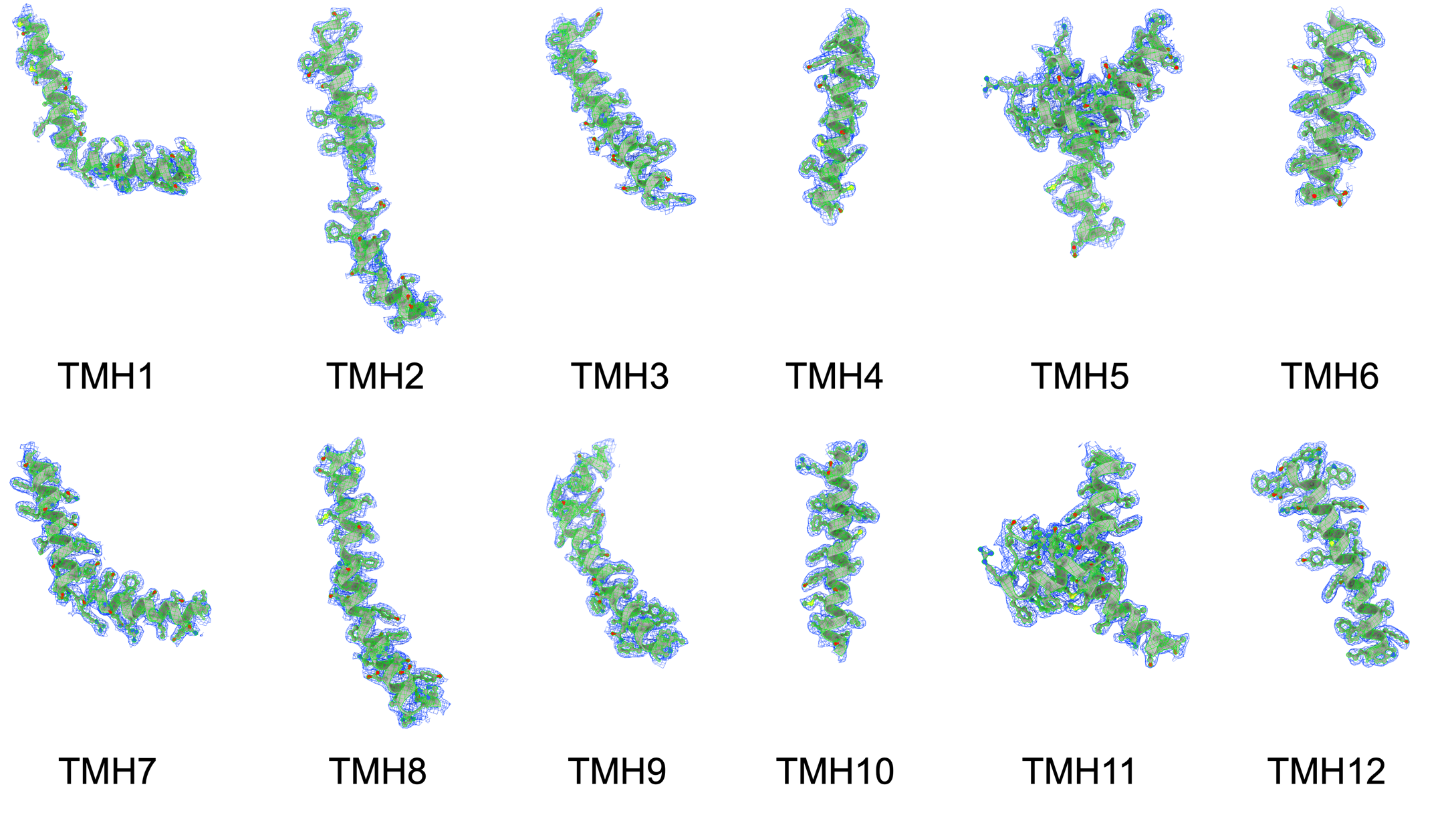

**Figure S2** Portions of the cryo-EM map showing the density around the transmembrane helices of Pdr5.

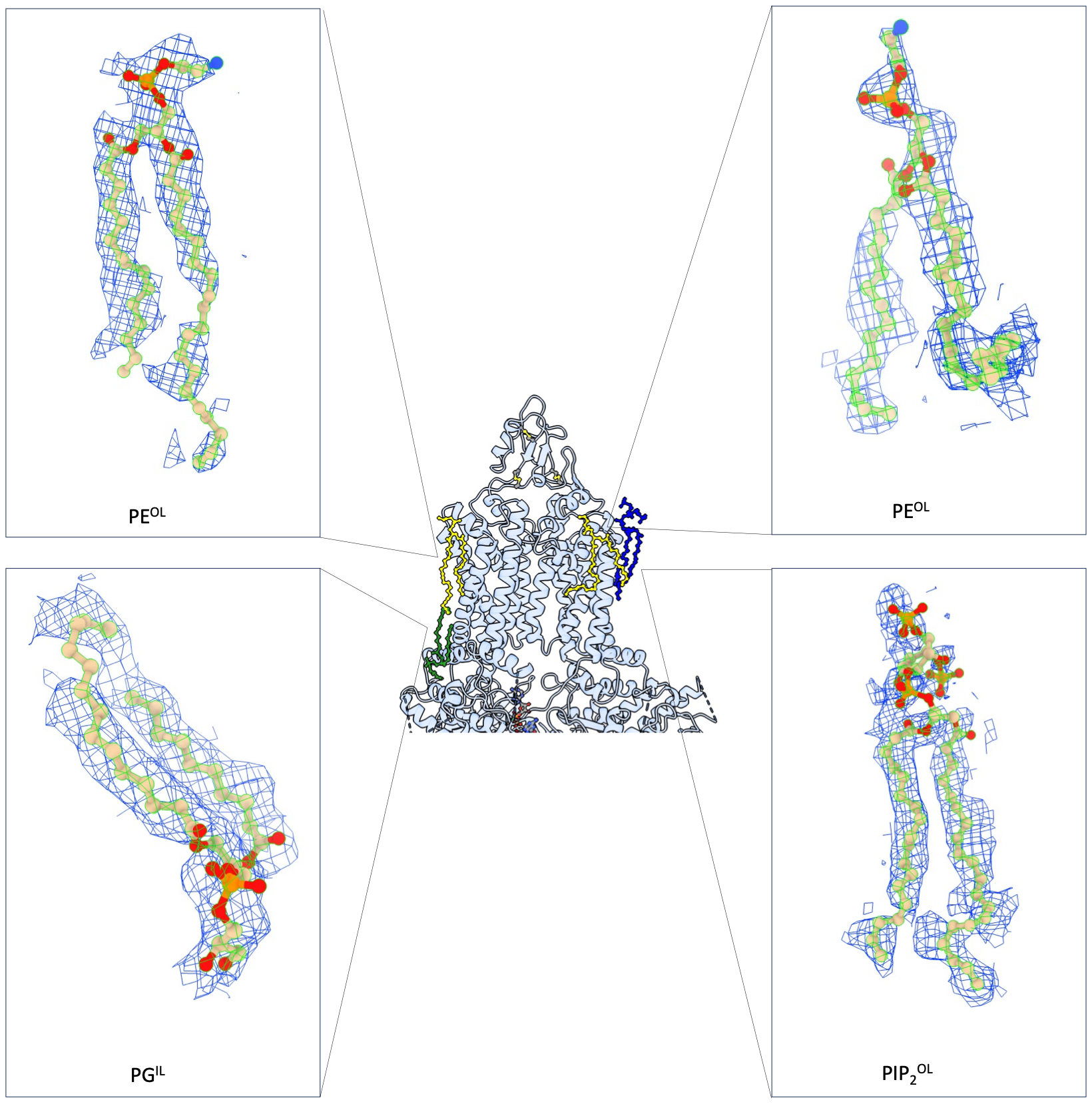

**Figure S3** Portions of the cryo-EM map showing the bound lipids in the Pdr5 structure: Phosphatidylinositol-4,5-bisphosphate (PIP_2_) in the outer leaflet (OL), phosphatidylglycerol (PG) in the inner leaflet, and two phosphatidylethanolamine (PE)) molecules in the outer leaflet of the phospholipid bilayer.

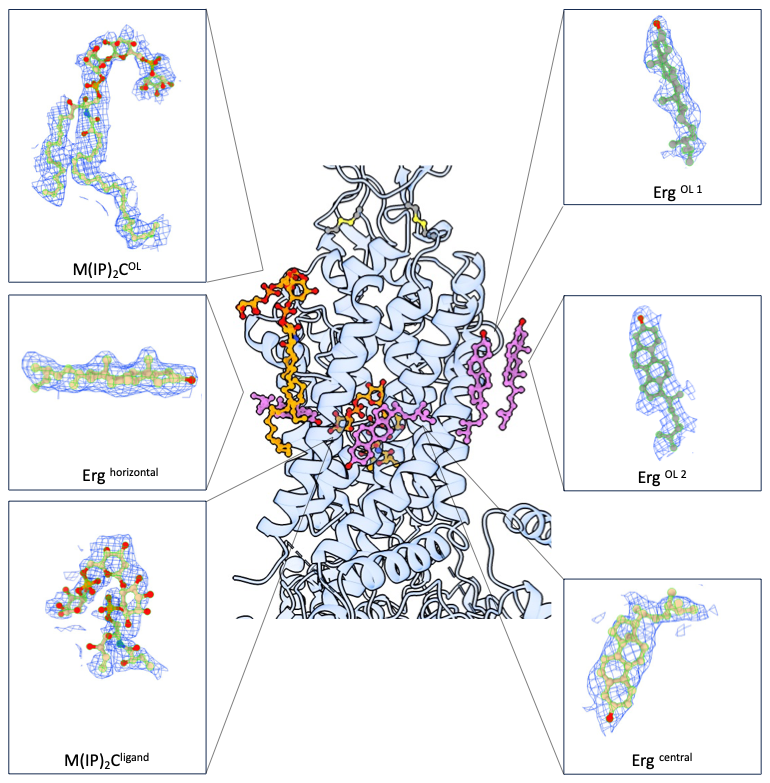

**Figure S4** Portions of the cryo-EM map showing the bound sphingolipids and ergosterol in the Pdr5 structure.

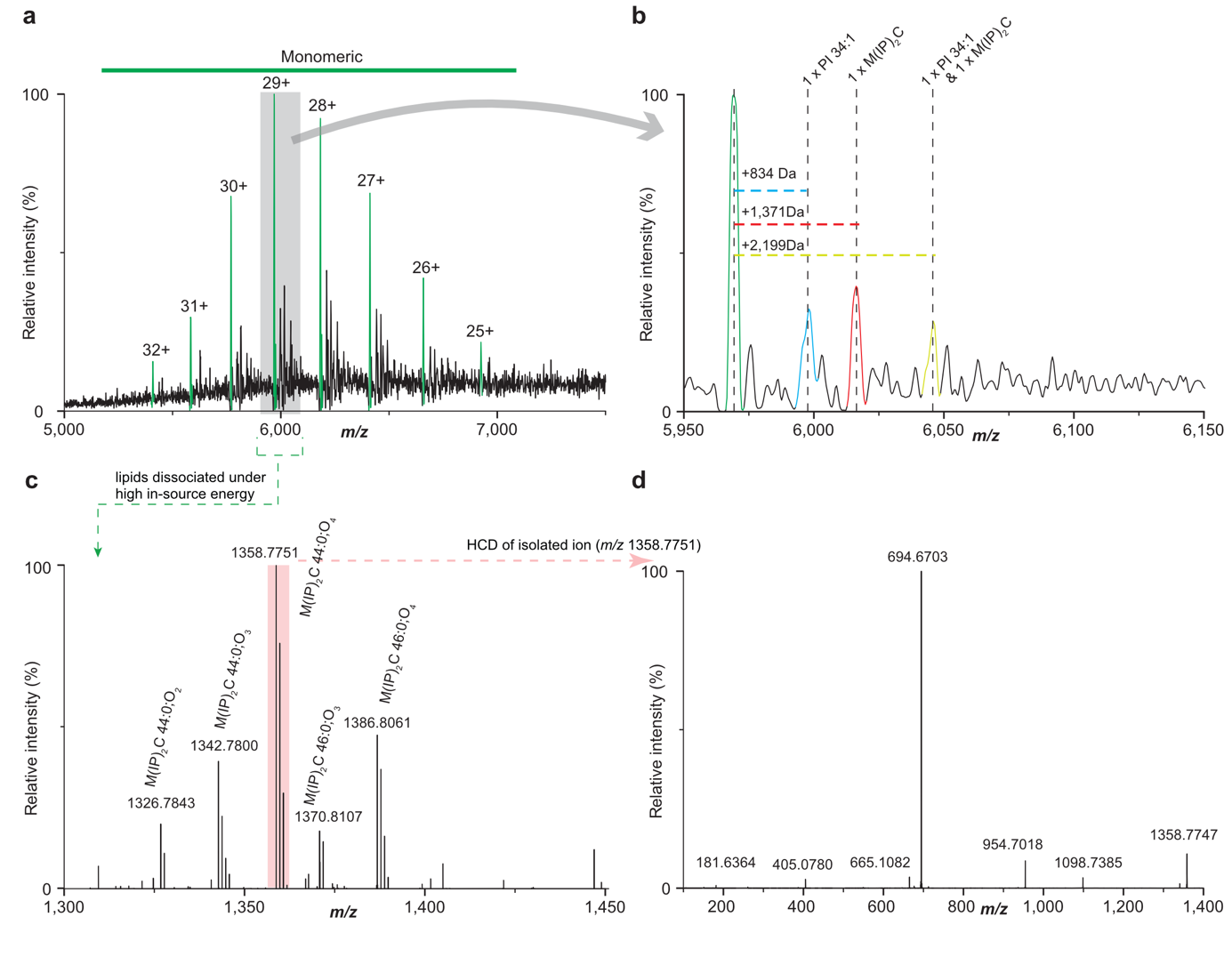

**Figure S5** Native mass spectrometry identifies Pdr5-bound M(IP)_2_C sphingolipid. (a) Native mass spectrometry (nMS) of purified Pdr5-N734A following λ-phosphatase treatment, showing the monomeric charge state distribution. (b). Expanded view of the 29+ charge state, showing satellite peaks with mass shift of +834 Da (PI 34:1), +1,371 Da (M(IP)_2_C) and +2,199 Da (PI 34:1 and M(IP)_2_C). (c) Pseudo-MS^2^ analysis of N734A-bound ligands in the *m/z* 1,300­­–1,450 range, revealing a series of M(IP)_2_C sphingolipid species. (d) Pseudo-MS^3^ analysis of the ion at *m/z* 1,358.7751 ion, confirming its identity as M(IP)_2_C 44:0;O4.

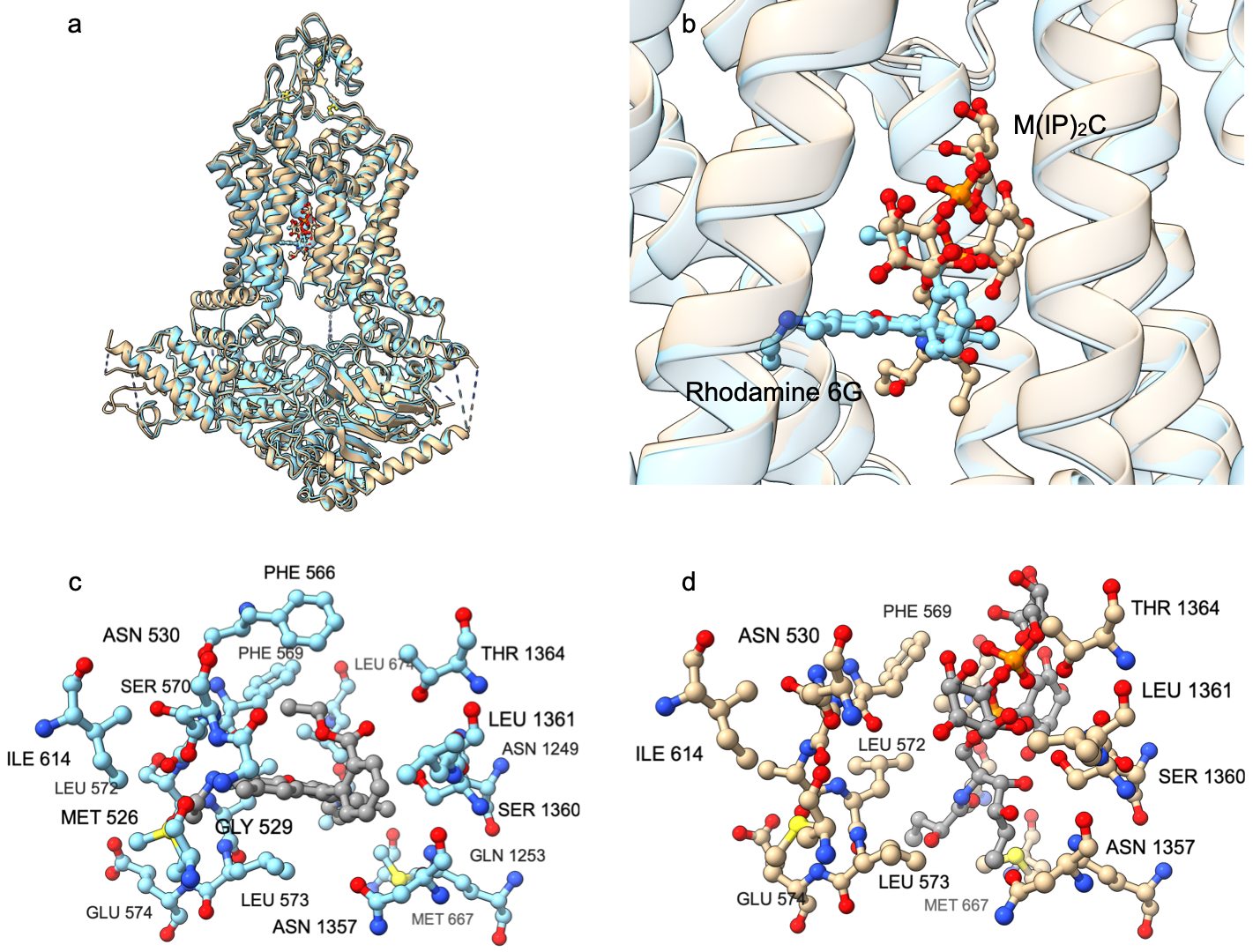

**Figure S6** (a) Superimposition of the inward-facing structures of Pdr5 in complex with M(IP)_2_C (light brown) and R6G (cyan) in the substrate-binding site with a rmsd value 0.78 Å over 1353 C_α_ atoms. (b) Zoom-in view of the substrate-binding site with the M(IP)_2_C-bound and R6G-associated structures shown as light brown and cyan cartoons, respectively. The two ligands are shown in ball-and-stick representation. Residues within a distance of 4 Å of R6G (c) and M(IP)_2_C (d) are shown in ball-and-stick representations and labeled.

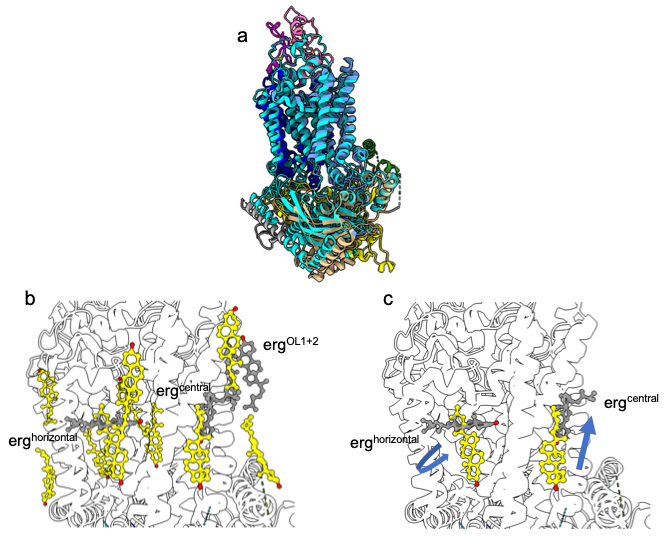

**Figure S7** (a) Superimposition of Pdr5 and ABCG2 (pdb entry 6HIJ)^1^ with an rmsd of 1.24 Å over 537 C_α_ atoms. Color-coding of Pdr5 is identical to Fig. 1a and ABCG2 is shown in cyan cartoon representation. (b) Location of cholesterol molecules bound to ABCG2 (yellow) and of ergosterol associated to Pdr5 (grey, labeled). Orientation is an in (a). (c) Comparison of erg^horizontal^ and erg^central^ (labeled) with their counterparts in ABCG2. Possible re-arrangements to achieve identical position are indicated by blue arrows.

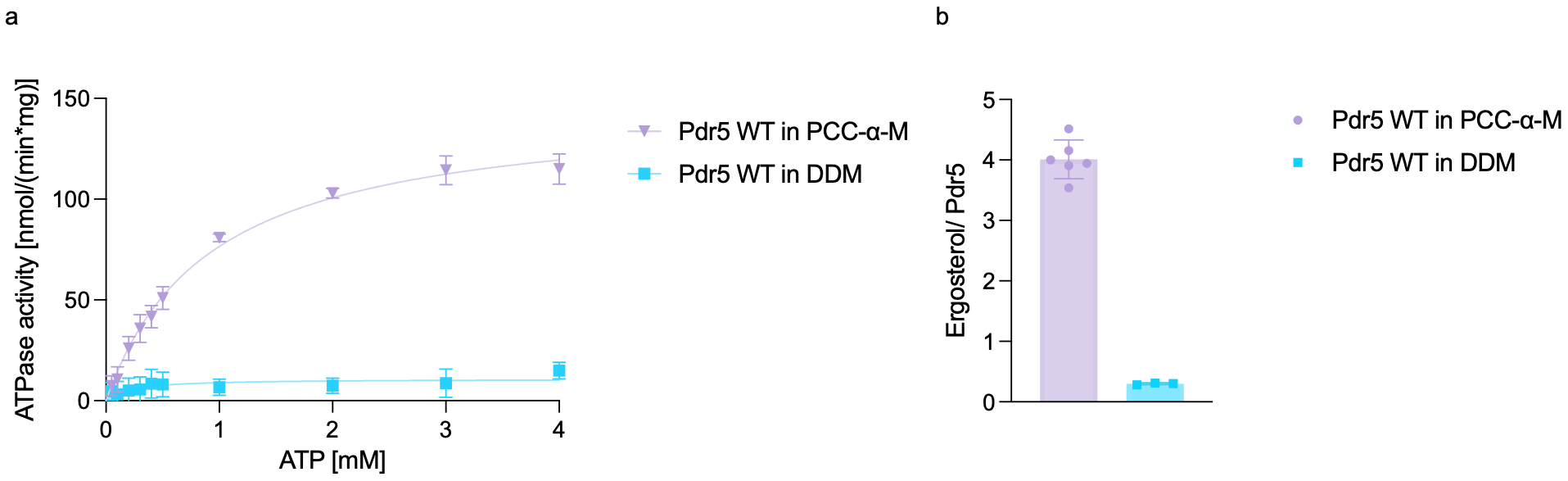

**Figure S8** (a) ATPase-activity of detergent-solubilized Pdr5 in PCC-α-M and DDM. (b) Ergosterol-quantification of detergent-solubilized Pdr5 in either PCC-α-M or DDM via GC-MS.

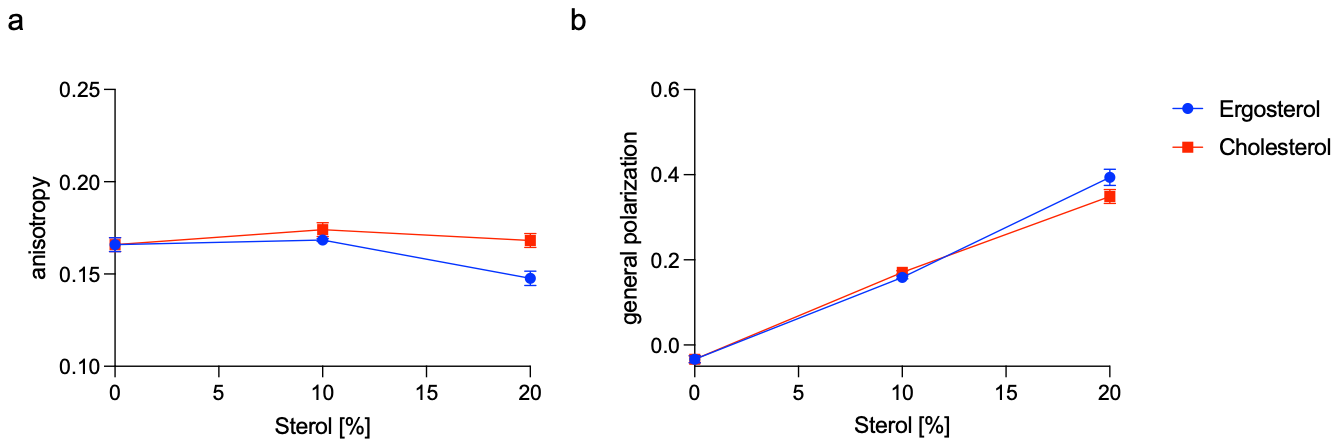

**Figure S9** Measurement of the membrane properties by insertion of the fluorescent probe laurdan into liposomes containing different amounts of cholesterol and ergosterol. Both the anisotropy (a) and the general polarization (b) of the liposomes were observed. Error bars represent the SD of at least three independent experiments

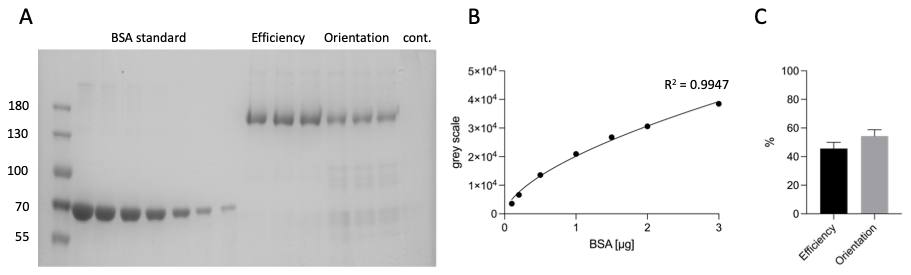

**Figure S10** Representative evaluation of the efficiency of the reconstitution and orientation of Pdr5 reconstituted into liposomes. The SDS-PAGE (a) shows a marker, a standard of bovine serum albumin (BSA standard) in the range of 3 to 0.1 µg, the samples determining the efficiency of the reconstitution (Efficiency) and orientation (Orientation) of Pdr5 reconstituted into liposomes (addition of trypsin), as well as a negative control (addition of trypsin and 0.1 % Triton-X-100: cont). (b) Protein amounts were quantified using a BSA standard curve based on Coomassie staining intensity. Due to the nonlinear staining response over the investigated concentration range, the standard curve was fitted using a power function ($y=a* x^{b}$) rather than linear regression ^2^. The fraction of reconstituted Pdr5 and outward-facing NBDs were calculated (c). Here, the signal intensities (‘Orientation’) were compared to the intensities of the ‘Efficiency’ signals indicating the amount of protein that was not digested by the trypsin. This amount represents Pdr5 reconstituted in an orientation in which the NBD face the external space. Error bars represent the SD of at least three independent experiments.

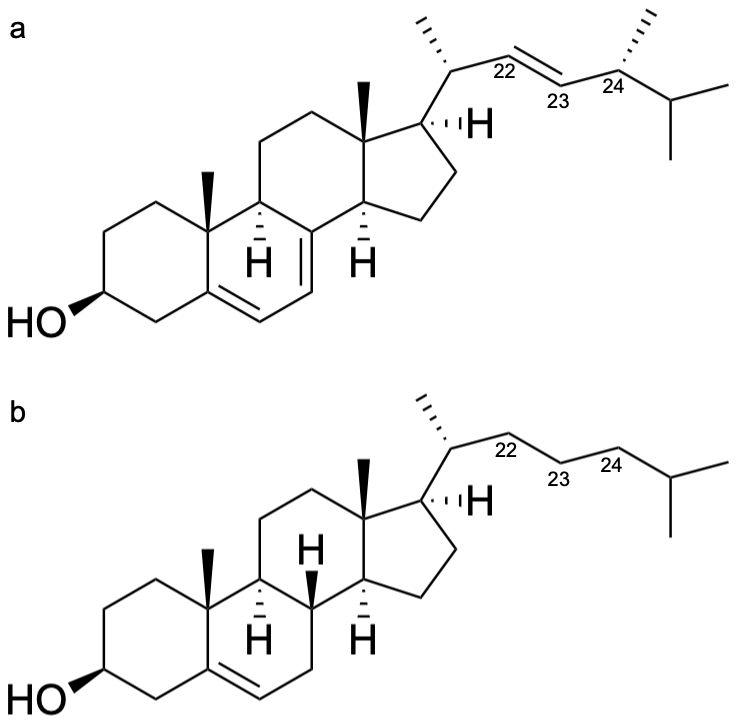

**Figure S11** Structural comparison of ergosterol and cholesterol. Chemical structure of (a) ergosterol and (b) cholesterol. Carbon atoms C22–C24 are labeled to highlight the structural differences between the two sterols, namely the C22–C23 double bond and the additional methyl group at C24 in ergosterol. Structures were generated using ChemDraw.

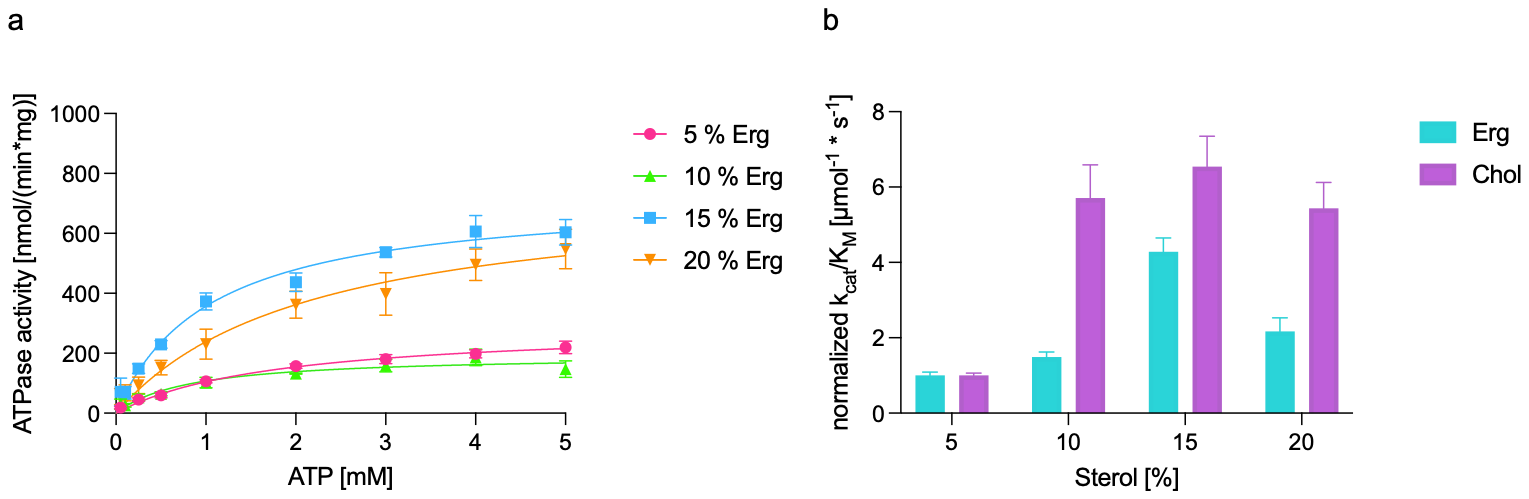

**Figure S12** ATPase activity of Pdr5 L/A-Y/A reconstituted in synthetic liposomes in the presence of varying amounts of ergosterol (a) and corresponding catalytic efficiency normalized to either 5 % (w/w) ergosterol or cholesterol, respectively (b). Error bars represent the SD of at least three independent experiments.

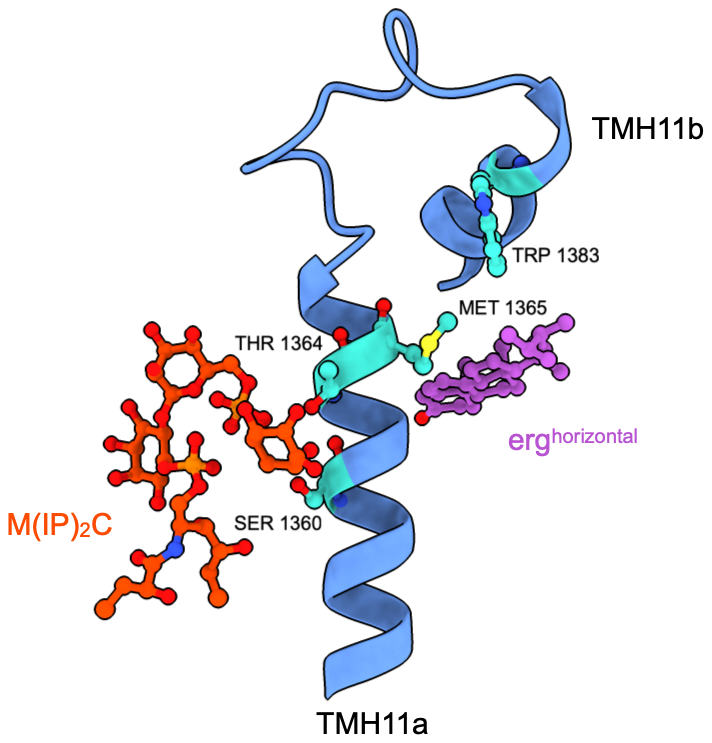

**Figure S13** Within the extended network of Pdr5-mediated interactions between the two M(IP)_2_C molecules and the two ergosterol molecules, erg^central^ and erg^horizontal^, respectively, four residues of Pdr5 have been functionally characterized previously. Focus on transmembrane helix 11a and b shown as blue cartoon connecting the M(IP)_2_C located in the substrate-binding site (shown in orange ball-and-stick representation) and erg^horizontal^ (shown in magenta ball-and-stick representation). Previously investigated residues S1360 and T1364 in TMH11a and M1365 and W1383 in TMH11b are shown in cyan ball-and-stick representation.

**Supplementary Tables**

**Supplementary Table 1** Cryo-EM data collection and refinement statistics

|  | **Pdr5 ATP/ADP**  **(EMDB-100182)**  **(PDB 34VA)** |
| --- | --- |
| **Data collection and processing** |  |
| Magnification | 105,000 |
| Voltage (kV) | 300 |
| Electron exposure (e^–^/Å^2^) | 48.9 |
| Defocus range (μm) | 0.7–2.2 |
| Pixel size (Å) | 0.831 |
| Symmetry imposed | C1 |
| Initial particle images (no.) | 632,196 |
| Final particle images (no.) | 124,340 |
| Map resolution (Å)  FSC threshold | 2.2*  0.143 |
| Map resolution range (Å) | 1.6- 5.5 |
| **Refinement** |  |
| Initial model used (PDB code) | 7P04 |
| Model resolution (Å)  FSC threshold | 2.2  0.143 |
| Map sharpening *B* factor (Å^2^) | –45.4 |
| Model composition  Non-hydrogen atoms  Protein residues  Ligands | 12560  1394  13 |
| *B* factors (Å^2^)  Protein  Ligand  Water | 76.3  105.7  80.9 |
| R.m.s. deviations  Bond lengths (Å)  Bond angles (°) | 0.009  1.058 |
| Validation  MolProbity score  Clashscore  Poor rotamers (%) | 1.37  5.37  1.0 |
| Ramachandran plot  Favored (%)  Allowed (%)  Disallowed (%) | 97.6  2.4  0 |

*: after density modification of the initial map (resolution of 2.45 Å).

**Supplementary Table 2** Kinetic parameters of the ATPase activity of Pdr5 wild type and mutants reconstituted in synthetic liposomes. Errors represent standard deviations of at least three independent experiments.

| Pdr5 | Erg [wt%] | Chol [wt%] | K_M_ [µM] | V_max_ [nmol/mg*min] | k_cat_/K_M_ [1/s*µM] | Difference with 5 % (w/w) sterol k_cat_/K_M_ |
| --- | --- | --- | --- | --- | --- | --- |
| Wild type | 5 | – | 1.12 ± 0.37 | 206 ± 23 | 0.66 ± 0.49 | – |
|  | 10 | – | 2.81 ± 1.09 | 269 ± 49 | 0.33 ± 0.21 | ns |
|  | 15 | – | 0.85 ± 0.17 | 642 ± 39 | 2.49 ± 0.54 | ** |
|  | 20 | – | 1.26 ± 0.33 | 420 ± 38 | 1.13 ± 0.72 | ns |
|  | – | 5 | 1.09 ± 0.09 | 1026 ± 28 | 2.77 ± 0.63 | – |
|  | – | 10 | 1.02 ± 0.20 | 943 ± 58 | 2.02 ± 1.08 | ns |
|  | – | 15 | 1.55 ± 0.24 | 1046 ± 59 | 1.72 ± 0.44 | ns |
|  | – | 20 | 1.28 ± 0.17 | 927 ± 42 | 2.08 ± 0.44 | ns |
| L1329A-Y1478A | 5 | – | 1.80 ± 0.24 | 293 ± 16 | 0.47 ± 0.04 | – |
|  | 10 | – | 0.82 ± 0.19 | 195 ± 13 | 0.69 ± 0.06 | ns |
|  | 15 | – | 1.05 ± 0.12 | 730 ± 28 | 1.99 ± 0.17 | **** |
|  | 20 | – | 2.15 ± 0.46 | 752 ± 68 | 1.01 ± 0.17 | * |
|  | – | 5 | 2.32 ± 0.35 | 659 ± 44 | 0.81 ± 0.05 | – |
|  | – | 10 | 0.36 ± 0.09 | 825 ± 46 | 4.61 ± 0.71 | ** |
|  | – | 15 | 1.03 ± 0.22 | 1956 ± 134 | 5.28 ± 0.65 | *** |
|  | – | 20 | 0.68 ± 0.12 | 1186 ± 56 | 4.39 ± 0.55 | ** |
| Pdr5 E1036Q | 15 | – | 0.22 ± 0.12 | 64 ± 7 | – | – |
|  | – | 15 | 0.38 ± 0.24 | 126 ± 17 | – | – |

Note: Analysis was carried out using a one-way ANOVA test in Prism 11.0.2 (GraphPad Inc.). ****: p < 0.0001, ***: p < 0.0002, **: p < 0.0021, *: p < 0.0332 ns: p < 0.1234.

1.	Jackson, S. M.*, et al.* Structural basis of small-molecule inhibition of human multidrug transporter ABCG2. *Nature Structural & Molecular Biology* **25**, 333-340 <https://doi.org/10.1038/s41594-018-0049-1> (2018).

2.	Splittgerber, A. G. & Sohl, J. Nonlinearity in protein assays by the Coomassie blue dye-binding method. *Anal Biochem* **179**, 198-201 <https://doi.org/10.1016/0003-2697(89)90225-x> (1989).
